# *Yap* haploinsufficiency leads to Müller cell dysfunction and late-onset cone dystrophy

**DOI:** 10.1101/2020.02.18.953943

**Authors:** Christel Masson, Diana García-García, Juliette Bitard, Élodie-Kim Grellier, Jérôme E. Roger, Muriel Perron

## Abstract

Hippo signalling regulates eye growth during embryogenesis through its effectors YAP and TAZ. Taking advantage of a *Yap* heterozygous mouse line, we here sought to examine its function in adult neural retina, where YAP expression is restricted to Müller glia. We first discovered an unexpected temporal dynamic of gene compensation. At post-natal stages, *Taz* upregulation occurs, leading to a gain of function-like phenotype characterized by EGFR signalling potentiation and delayed cell cycle exit of retinal progenitors. In contrast, *Yap^+/-^* adult retinas no longer exhibit TAZ-dependent dosage compensation. In this context, *Yap* haploinsufficiency in aged individuals results in Müller glia dysfunction, late-onset cone degeneration and reduced cone-mediated visual response. Alteration of glial homeostasis and altered patterns of cone opsins were also observed in Müller cell specific conditional *Yap* knockout mice. Together, this study highlights a novel YAP function in Müller cells for the maintenance of retinal tissue homeostasis and the preservation of cone integrity. It also suggests that YAP haploinsufficiency should be considered and explored as a cause of cone dystrophies in human.

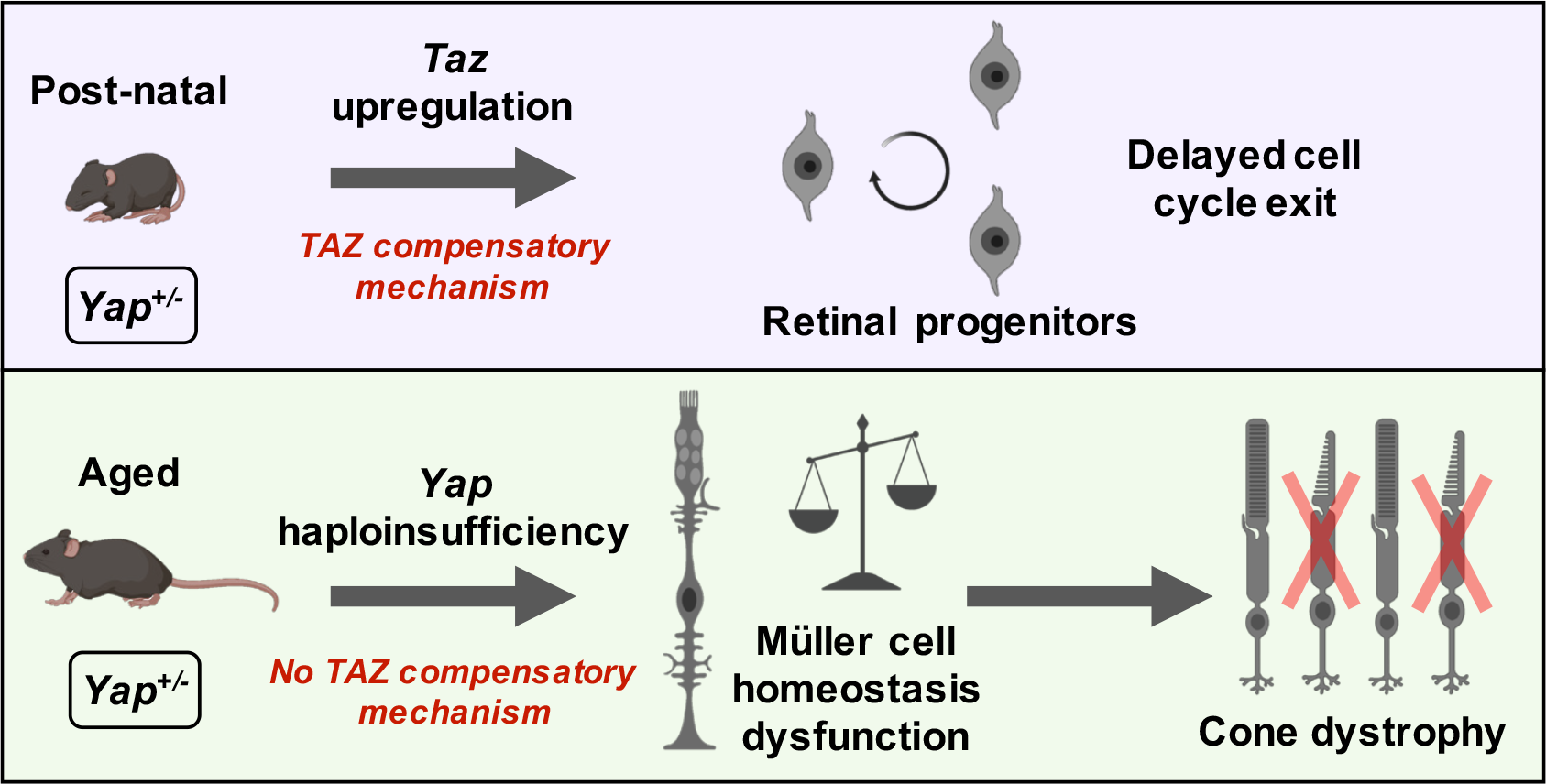

**Main Points:**

- This study finds unsuspected dynamics of YAP compensatory mechanisms in the retina of *Yap^+/-^* mice
- *Yap* haploinsufficiency results in Müller glia dysfunction in aged mice, leading to late-onset cone dystrophy

## INTRODUCTION

Deregulation of key signalling pathways, such as Wnt or Notch, well known for their involvement in retinal development, are at the origin of retinal diseases in adults (Lad et al., 2009; Zheng et al., 2010). Studying developmental signalling pathways can thus allow the identification of possible causes of retinal disorders in adults, pinpointing possible targets for therapeutic approaches. The Hippo signalling tightly regulates eye development, and its deregulation in animal models leads to severe ocular defects (reviewed in (Lee et al., 2018; Moon and Kim, 2018)). This signalling pathway consists of a kinase cascade that ultimately phosphorylates the transcription coactivators YAP (Yes associated protein) and TAZ (WW domain–containing transcription regulator 1), causing their retention in the cytoplasm or their degradation. When the pathway is inactive, YAP/TAZ are translocated to the nucleus, leading to activation of their target genes (Edgar, 2006; Pan, 2007). During retinal development, YAP and TAZ are expressed in all optic vesicle compartments, becoming more prominently expressed in the retinal pigment epithelium (RPE) and the ciliary margin of the optic cup (Miesfeld et al., 2015; Moon and Kim, 2018). Studies in zebrafish and mice indicate that YAP is necessary for the maintenance of progenitor populations in both the retina and the RPE (Jiang et al., 2009; Zhang et al., 2012; Asaoka et al., 2014; Miesfeld et al., 2015; Kim et al., 2016; Moon and Kim, 2018). Besides, mutations in Hippo pathway components have been identified in human ocular diseases, such as the Sveinsson’s chorioretinal atrophy (SCRA) (Fossdal et al., 2004; Kitagawa, 2007), in patients with ocular manifestations of the neurofibromatosis type-2 human disease (Asthagiri et al., 2009), or in patients exhibiting coloboma (Williamson et al., 2014; Oatts et al., 2017). Together, these studies have contributed to highlight the important role of the Hippo-YAP pathway in eye development and ocular disorders. Although it has previously been established that YAP remains expressed in the adult retina in both Müller glia and RPE cells (Hamon et al., 2017), its function in the adult eye has received little attention thus far.

To investigate YAP function in the adult, and because *Yap* knockout is lethal at embryonic day 8.5 (Morin-kensicki et al., 2006), we took advantage of a *Yap* heterozygous mouse model. We found that depletion of one *Yap* allele leads to a transient TAZ compensatory mechanism at post-natal stages, associated with an increased expression of YAP/TAZ target genes. Consistent with the known critical role of the Hippo pathway in regulating neural progenitor cell proliferation (Cao et al., 2008), we found that such enhanced TAZ activity in *Yap^+/-^* postnatal retinas is associated with a gain of function-like phenotype, *i.e.* delayed cell cycle exit of retinal progenitors. We identified the regulation of the EGFR (Epidermal Growth Factor Receptor) pathway as a potential underlying mechanism. Conversely, TAZ compensatory regulation declined during aging in *Yap* heterozygous retinas. We found that non-compensated decrease of *Yap* expression in aged retinas leads to altered Müller cell homeostatic function, causing late-onset cone degeneration and thereby impaired cone visual function. This work thus uncovers a novel unexpected role for YAP in cone photoreceptor maintenance and proper vision.

## METHODS AND MATERIAL

### Ethics statement

All animal experiments have been carried out in accordance with the European Communities Council Directive of 22 September 2010 (2010/63/EEC). All animal care and experimentation were conducted in accordance with institutional guidelines, under the institutional license D 91-272-105. The study protocols were approved by the institutional animal care committee CEEA n°59 and received an authorization by the “Ministère de l’Education Nationale, de l’Enseignement Supérieur et de la Recherche” under the reference APAFIS#1018-2016072611404304 v1.

### Mice

Mice were kept at 21°C, under a 12-hour light/12-hour dark cycle, with food and water supplied *ad libitum*. Heterozygous *Yap^+/-^* mice were obtained from Sigolene Meilhac (Institut Pasteur, Paris). Briefly, *Yap^flox/+^* mice (Reginensi et al., 2013) were crossed with PKG Cre mice (PGK-Cre transgene is maternally expressed and serves as a tool for early and uniform activation of the Cre site-specific recombinase (Lallemand et al., 1998)) to generate the *Yap^+/-^* mice, that are viable and fertile. *Yap^flox/flox^;Rax-CreER^T2^* mice were obtained as previously described (Hamon et al., 2019) by mating *Yap^flox/flox^* mice (Reginensi et al., 2013) with heterozygous Rax-CreER^T2^ knock-in mice (Pak et al., 2014). The Cre activity was induced through a single intraperitoneal injection of 4-hydroxy-t-moxifen (4-OHT; 1 mg/kg) at P10. Genotyping was done by PCR using genomic DNA prepared from mouse tail snips. Primers are provided in Supplementary Table S1.

### Tissue sectioning, immunohistochemistry, and EdU labelling

Eyes of sacrificed animals were rapidly enucleated and dissected in Hanks’ Balanced Salt solution (Gibco) to obtain posterior segment eye-cups, which were then fixed in 4% paraformaldehyde, 1X PBS, for 1 hour at 4°C. Eye-cups were then dehydrated, embedded in paraffin and sectioned (7µm) with a Microm HM 340E microtome (Thermo Scientific). Hematoxylin-eosin (H&E) staining was performed according to the manufacturer’s instructions (BBC Biochemical). Stained sections were mounted with Eukitt (Polylabo). Standard immunohistochemistry techniques on paraffin sections were applied with the following specificities: antigen unmasking treatment was done in boiling heat-mediated antigen retrieval buffer (10mM sodium citrate, pH 6.0) for 20 min. Primary antibody was diluted in ready-to-use diluent (Dako). Primary and secondary antibodies are listed in Supplementary Table S2. Sections were counterstained with 1µg/ml DAPI (Thermo Fisher Scientific) and mounted with Fluor Save^TM^ reagent (Millipore). For retinal flat mounts, small dorsal incisions were made before fixation to mark the orientation of the retina within the eyecup. For EdU incorporation experiments, mice were given a single 50 to 100 µl intra-peritoneal injection of 10 µM of EdU (Invitrogen) at the indicated stage. EdU incorporation was detected on flat mounted retinal mouse explants using the Click-iT EdU Imaging Kit (Invitrogen) according to manufacturer’s recommendations. For double labelling, EdU labelling was done first, followed by immunostaining.

### Imaging

Fluorescence images were acquired using a LSM710 confocal microscope (Zeiss). Whole retina images were acquired using an AXIOZoom.V16 (Zeiss) using the mosaic mode. Image mosaics of flat mounted retinas were acquired and combined by the stitching processing method using ZEN Tiles module (Zeiss). Brightfield images of H&E staining were acquired using an AxioImager.M2 microscope. Image processing was performed using Zen 2.1 (Zeiss), Fiji (National Institutes of Health, (Schindelin et al., 2012)) and Photoshop CS4 software (Adobe) software. The same magnification, laser intensity, gain and offset settings were used across animals for any given marker.

### Western blotting

Western blot was performed on protein extracts from single retinas, at least on 3 individuals per condition, unless otherwise specified in the figure legends. Retinas from enucleated eyes were dissected in Hanks’ Balanced Salt solution (Gibco) by removing the anterior segment, vitreous body, sclera and RPE and were frozen at -80°C. Retinas were lysed in P300 buffer (20 mM Na_2_HPO_4_; 250 mM NaCl; 30 mM NaPPi; 0.1% Nonidet P-40; 5 mM EDTA; 5mM DTT) supplemented with protease inhibitor cocktail (Sigma-Aldrich). For RPE protein extracts, the anterior segment and the retina were removed from enucleated mouse eyes. The RPE was then separated from the choroid by incubating the posterior eyecup (sclera-choroid-RPE) with P300 buffer for 10 min. Protein concentration was determined using a Lowry protein assay kit (DC Protein Assay; Bio-Rad). Equal amounts of proteins (20 µg/lane) of each sample were loaded, separated by 7.5% SDS-PAGE (Bio-Rad) and transferred onto nitrocellulose membranes. Western blots were then conducted using standard procedures. Primary and secondary antibodies are listed in Supplementary Table S2. An enhanced chemiluminescence kit (Bio-Rad) was used to detect the proteins. Each sample was probed once with anti-*α*-tubulin antibody for normalization. Quantification was done using Fiji software (National Institutes of Health, (Schindelin et al., 2012)).

### Electroretinographic analysis

Electroretinograms (ERGs) were recorded using a Micron IV focal ERG system (Phoenix Research Labs). Mice were dark-adapted overnight and prepared for recording in darkness under dim-red illumination. Mice were anesthetized with intraperitoneal injection of ketamine (90 mg/ kg, Merial) and xylazine (8 mg/kg, Bayer), and were topically administered tropicamide (0.5%) and phenylephrine (2.5%) for pupillary dilation. Flash ERG recordings were obtained from one eye. ERG responses were recorded using increasing light intensities ranging from -1.7 to 2.2 log cd.s/m^2^ under dark-adapted conditions, and from -0.5 to 2.8 log cd.s/m^2^ under a background light that saturates rod function. The interval between flashes varied from 0.7 s at the lowest stimulus strengths to 15 s at the highest ones. Five to thirty responses were averaged depending on flash intensity. Analysis of a-wave and b-wave amplitudes was performed using LabScribeERG software (https://www.iworx.com/research/software/labscribe). The a-wave amplitude was measured from the baseline to the negative peak and the b-wave was measured from the baseline to the maximum positive peak.

### Fluorescein Angiography

Mice were anesthetized as described above. Pupils were dilated using 0.5% tropicamide (Théa) and 5% chlorhydrate phenylephrine (Europhta) eye drops. The mouse was placed on the imaging platform of the Micron IV system (Phoenix Research Labs), and Ocry-gel was applied on both eye to keep the eye moist during the imaging procedure. Mice were injected in tail’s vein with 100 µL of 5% fluorescein isothiocyanate dextran (Sigma), and rapid acquisition of fluorescent images ensued for ∼5 minutes.

### RNA extraction and RT-qPCR

RT-qPCR experiment was performed on at least three mice per condition. Total RNA was extracted from a single retina using RNeasy mini kit (Qiagen) and treated with DNAse I according to the manufacturer’s instructions. RNA quantity was assessed using the NanoDrop 2000c UV-Vis spectrophotometer (Thermo Fisher Scientific). Total RNA (500 ng) was reverse transcribed in the presence of oligo-(dT)20 using Superscript II reagents (Thermo Fisher Scientific). For each RT-qPCR, cDNA was used in the presence of EvaGreen (Bio-Rad), and the reactions were performed in triplicates on a CFX96 Real-Time PCR Detection System (Bio-Rad). Differential expression analysis was performed using the ΔΔCt method and normalized using the geometric mean expression of two housekeeping gene, *Rps26* and *Srp72* (Vandesompele et al., 2002). For each gene, the relative expression in each sample was calculated using the mean of the controls as the reference (1 a.u.). Primers used are listed in Supplementary Table S1.

### Quantification and statistical analysis

The numbers of labelled cells on retinal section or retinal flat mounts were manually counted in a defined field (sizes are indicated in the legends). For all such counting, as well as qPCR and western blot analysis, at least three retinas were used per condition. The non-parametric Mann-Whitney test was used for all statistical analysis except for the ERG experiment for which a two-way *ANOVA* followed by *Bonferroni’s* multiple comparison test was used. All results are reported as mean ± SEM. All analyses were performed using GraphPad Prism 5.01 (GraphPad Software, La Jolla California USA) with statistical significance set at p-value*≤*0.05.

## RESULTS

### Compensatory *Taz* regulation in *Yap^+/-^* post-natal mice

In contrast with the developmental arrest observed in *Yap^-/-^* mice (Morin-kensicki et al., 2006), *Yap* heterozygous mice are viable and fertile. We wondered whether *Yap* gene haploinsufficiency could occur in the adult, and therefore decided to examine, in detail, the molecular and phenotypic consequences of one *Yap* allele deletion. As expected, qPCR and western blot analysis confirmed an approximately two-fold decrease of *Yap* expression in post-natal and adult *Yap^+/-^* retinas compared to controls (Figure 1A, B). Noticeably, the decrease appeared more severe during aging. By immuno-histochemical (IHC) analysis, we found that YAP expression was severely diminished in both Müller glial and RPE cells, where YAP has previously been shown to be expressed (Hamon et al., 2017) (Figure 1C). Prior to analysing potential ocular phenotypes in *Yap^+/-^* mice, we wanted to confirm that the observed *Yap* expression reduction actually leads to the decrease of *Yap* target gene expression. Indeed, this may not be the case since compensatory regulation mechanisms have previously been reported between YAP and TAZ (Miesfeld et al., 2015; Deng et al., 2017; Neto et al., 2018). In accordance with this, we observed an increase in *Taz* transcript abundance in post-natal *Yap^+/-^* retinas compared to controls (Figure 1D). Noticeably, the compensatory upregulation waned in adult stages (after P21). This trend was confirmed at the protein level (Figure 1E). In order to assess the net outcome of the compensatory mechanism on YAP/TAZ activity, we analysed by RT-qPCR the expression level of *Ctgf* and *Cyr61*, which have been recognized as direct YAP/TAZ target genes (Lai et al., 2011). Similar to *Taz* expression profile, we found that both genes were first upregulated at post-natal stages (prior P21), but downregulated from 8-month onwards in *Yap^+/-^* mice compared to controls (Figure 1D). YAP/TAZ interacting transcription factors TEAD1 and TEAD2 followed the same expression profile (Figure 1D and 1E). Of note, TEAD3 and TEAD4 are not expressed in the mouse retina (Hamon et al., 2017). Together, these data reveal the existence of a dynamic regulatory mechanism taking place in *Yap^+/-^* mice, that first leads to an increased expression of key YAP related and target genes at post-natal stages due to the compensatory overexpression of TAZ that diminished progressively with aging, with a concomitant decreased expression of these genes in adult mice. This unexpected temporally dynamic compensation prompted us to undertake a thorough phenotypic analysis of *Yap^+/^*^-^ mice retina at both post-natal and adult stages, when compensatory mechanisms are occurring or not, respectively.

**Figure 1.**
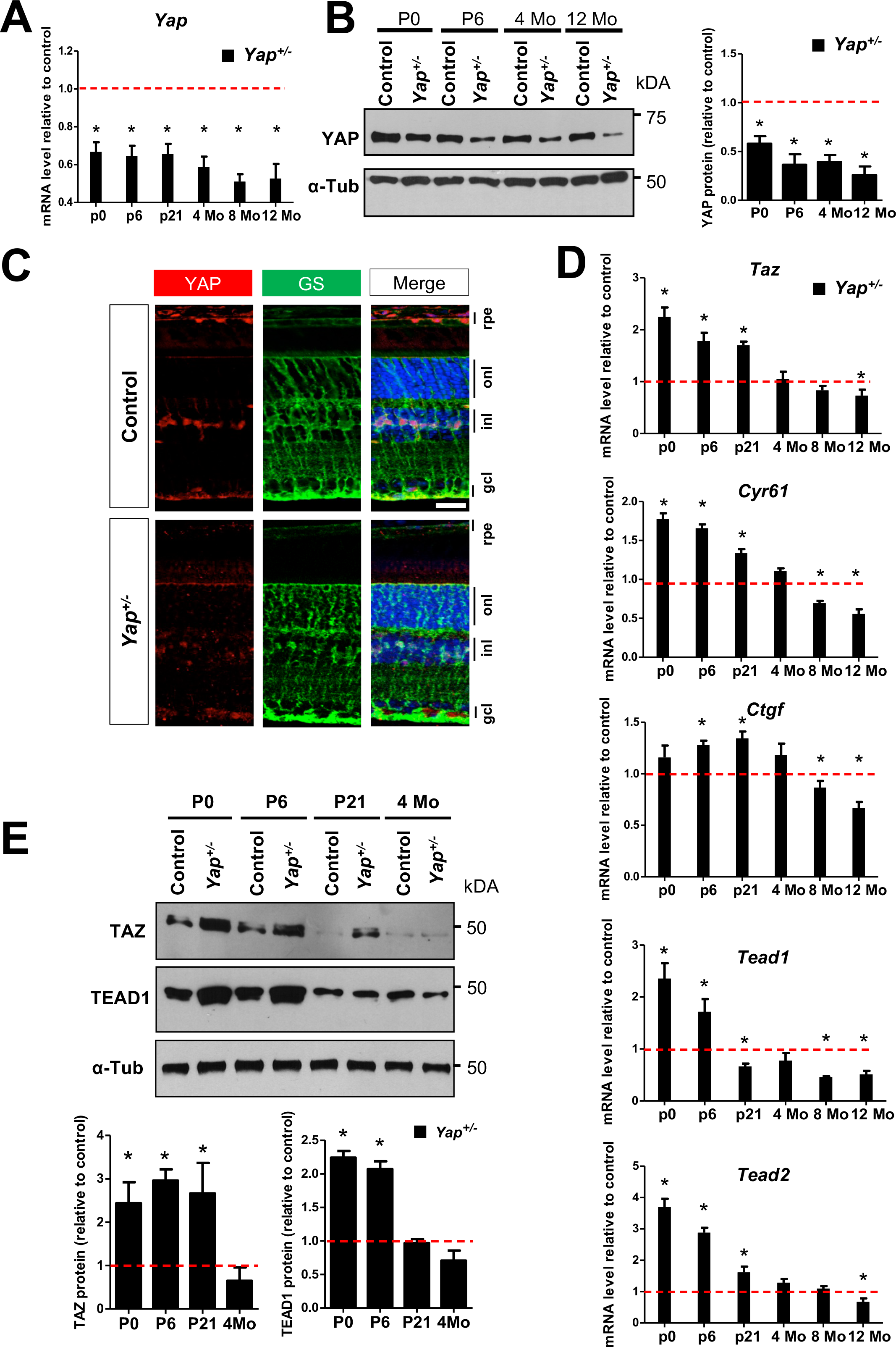
Compensatory regulation in *Yap*^+/-^ mice. (**A**) RT-qPCR analysis of *Yap* expression in *Yap^+/-^* mice retina, relative to wild type controls (dashed line) (n=3 biological replicates per condition). (**B**) Analysis of YAP protein expression level by western blot. The quantification is normalized to α-tubulin (α-Tub) signal and relative to wild type controls at each stage (dashed line) (n=3 retinas for each condition). (**C**) 4-month-old mice retinal sections immunostained for YAP (red) and a marker of Müller cells, glutamine synthetase (GS, green). Nuclei are DAPI counterstained (blue). (**D**) RT-qPCR analysis of *Taz*, *Cyr61*, *Ctgf*, *Tead1* and *Tead2* expression relative to wild type controls (dashed line) (at least 3 biological replicates per condition). (**E**) Western blots analysis of TAZ and TEAD1. The quantification is normalized to α-tubulin (α-Tub) signal and relative to controls at each stage (dashed line) (n=3 retinas for each condition). RPE: retinal pigment epithelium; INL: inner nuclear layer; ONL: outer nuclear layer; GCL: ganglion cell layer. All values are expressed as the mean ± SEM. Statistics: Mann-Whitney test, *p≤ 0.05. Scale bar: 20 µm.

### Delayed cell cycle exit of retinal progenitor cells in *Yap*^+/-^ post-natal mice

*Yap* overexpression in new-born mice was shown to promote retinal cell proliferation, while *Yap* knock-down leads to the opposite phenotype (Zhang et al., 2012). Therefore, considering the abundance changes of both YAP (decrease) and TAZ (increase) in *Yap* heterozygous postnatal retinas, we wondered how it impacts the proliferative behaviour of retinal progenitor cells. Interestingly, 24 hours after EdU intraperitoneal injection at P5 (when most progenitors have exited the cell cycle), four times more EdU-positive cells were detected in the central part of mutant retinas compared to control ones (Figure 2A, B). This persistence of a population of proliferative cells in *Yap*^+/-^ P6 retinas was supported by IHC analysis, showing more cells labelled with the cell proliferation markers PCNA and CyclinD1, compared to controls (Figure 2C). We found no more EdU-labelled cells in *Yap^+/-^* retinas, neither in the central nor in the peripheral region, when EdU injection was performed at P11 (Supplementary Figure S1), suggesting that *Yap^+/-^* late progenitors eventually exited the cell cycle between P6 and P11. We next sought to determine, by EdU birthdating, the fate of these late progenitor cells. Proliferative progenitors in P6 *Yap^+/-^* retinas gave rise to both late born neurons (CHX10-positive bipolar cells and Recoverin-positive photoreceptors) and glial cells (SOX9-positive Müller cells) in the central retina (Supplementary Figure S2A-D). To be able to compare this distribution with a control one, and since no EdU-positive cells were present in the central retina of P6 control individuals, we analysed the fate of EdU cells localized in the periphery of the retina. The proportion of double EdU/CHX10-, EdU/Recoverin- and EdU/SOX9-positive cells among EdU cells was similar in *Yap^+/-^* and in control retinas (Supplementary Figure S2E). Finally, a series of IHC staining on P21 animals did not reveal any significant differences between wild type and *Yap^+/-^* retina regarding markers of rods, cones, bipolar, ganglion, amacrine, horizontal, and Müller cells (data not shown). Together, these data suggest that a subset of *Yap^+/-^* retinal progenitor cells have a delayed cell cycle exit, without exhibiting any apparent bias in their cell fate determination.

**Figure 2.**
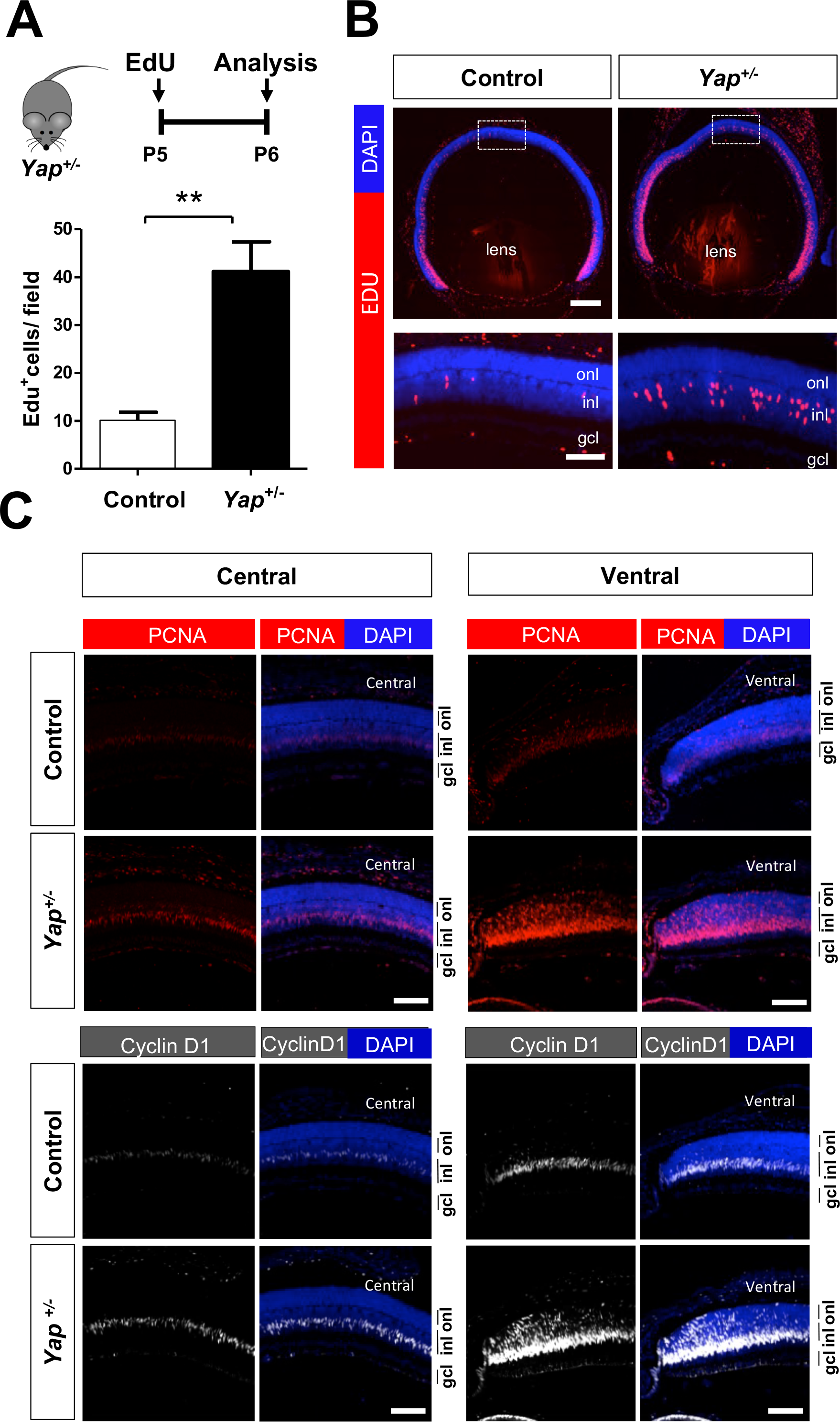
Prolonged proliferation of retinal progenitors at postnatal stages in *Yap^+/-^* mice. (A) Timeline diagram of the experimental procedure used in B. Wild-type (Control) or *Yap^+/-^* mice were injected with EdU at P5 and analyzed 24 hours later. (**B**) P6 retinal sections labelled for EdU (red) and stained with DAPI (blue). The delineated areas are enlarged in the bottom panels. Histogram represents the number of EdU^+^ cells per field (250 µm x 250 µm). Values ± SEM from 6 wild type control retinas and 5 *Yap^+/-^* retinas are shown. (**C**) P6 retinal sections immunostained for PCNA (red) or Cyclin D1 (grey). Both central and peripheral ventral regions are shown. Nuclei are DAPI counterstained (blue). INL: inner nuclear layer; ONL: outer nuclear layer; GCL: ganglion cell layer. Statistics: Mann-Whitney test, **p≤ 0.01. Scale bar: 200 µm (B) and 50 µm (C and enlarged panels in B).

### EGFR pathway potentiation in *Yap^+/-^* post-natal mice

YAP has been recognized as an integrator of several key signalling pathways. In particular, crosstalk between EGFR and Hippo pathways have been reported (He et al., 2015; Chen and Harris, 2016; Yang et al., 2016; Hamon et al., 2019). Knowing that EGFR signalling regulates proliferation of retinal progenitor cells at postnatal stages (Close et al., 2006), we hypothesized that the excess of late retinal proliferative progenitor cells in *Yap^+/-^* retinas could result from a deregulated EGFR pathway. RT-qPCR analysis at P0 and P6 showed a significant upregulation of *Egfr*, *Erbb2, Erbb3* and *Erbb4* (encoding receptors of the EGFR family: EGFR, Her2, Her3 and Her4, respectively), as well as of *Hbegf* and *Neuregulin 1* (encoding EGFR ligands: HB-EGF and NRG1, respectively), in *Yap^+/-^* mice compared to controls (Figure 3A). From P21, their expression diminished progressively in *Yap^+/-^* mice to reach control levels or levels lower than the controls (Figure 3A). It is established that EGFR signalling acts through the activation of the MAPK, PI3K/AKT and STAT3 pathways and previous reports have shown that these pathways are activated during proliferation of retinal progenitors cells (Zhang et al., 2005; Oliveira et al., 2008; Ornelas et al., 2013). We thus investigated whether their activity was affected in *Yap*^+/-^ mice. P-AKT/AKT and P-STAT3/STAT3 ratios were increased in *Yap*^+/-^ mice compared to controls at P6, while no difference was observed for P-ERK/ERK ratio (Figure 3B). Together, these results revealed that EGFR pathway activity is potentiated at early postnatal stages in *Yap^+/-^* retina, and that this upregulation no longer occurs in adult stages. We therefore propose that the TAZ-dependent compensatory regulation that we observed in *Yap^+/-^* mice during post-natal stages may underlie this EGFR pathway activation.

**Figure 3.**
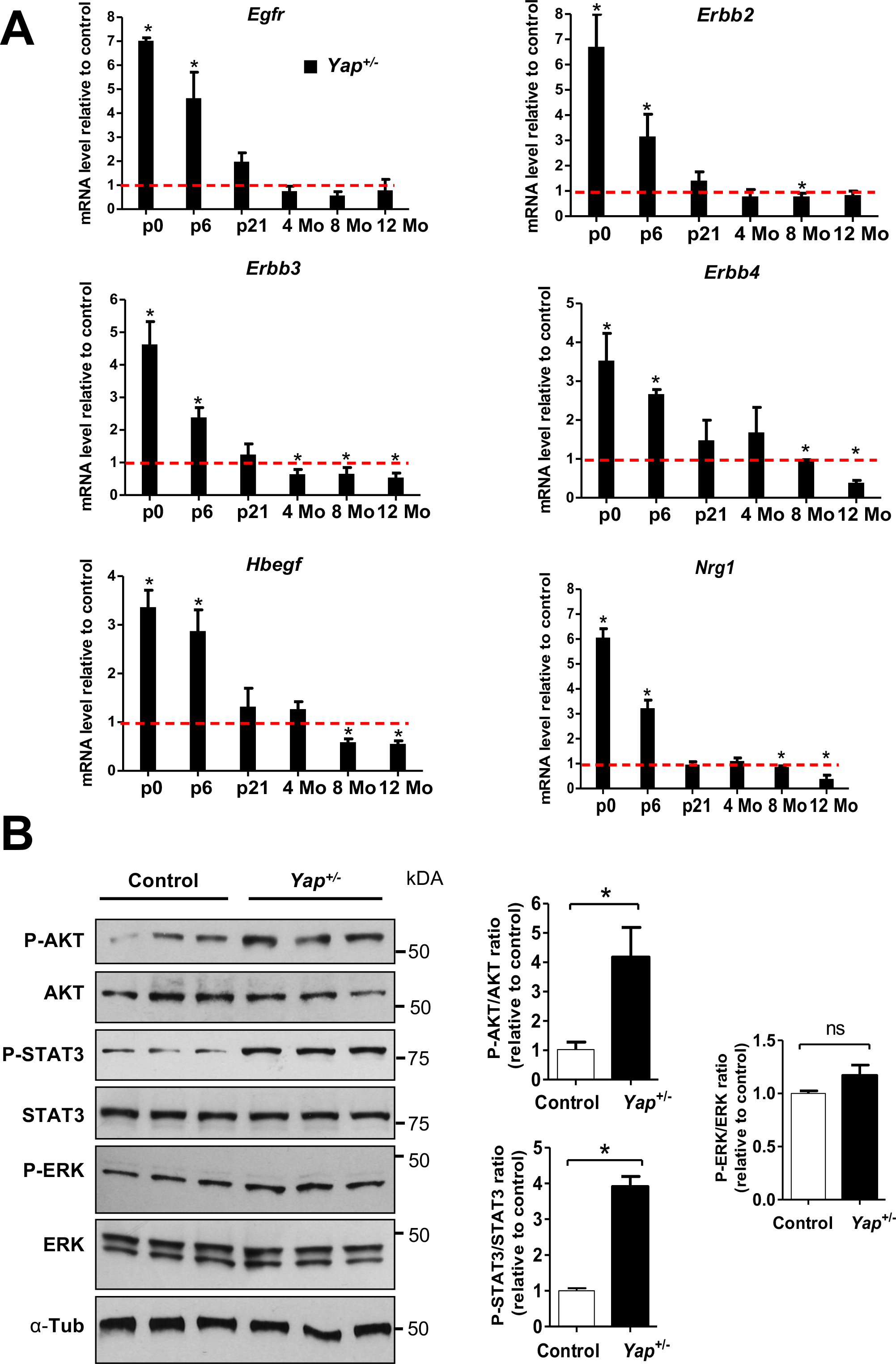
EGFR pathway activation in the retina of *Yap^+/-^* post-natal mice. (**A**) RT-qPCR analysis of various EGFR pathway signaling gene expression (*Egfr, Erbb2, Erbb3, Hbegf, Nrg1*), relative to wild type controls (dashed lines) (at least 3 biological replicates per condition). (**B**) Analysis of protein expression levels of EGFR signaling pathway components at P6 by western blot. The quantification of p-AKT/AKT, p-STAT3/STAT3 and p-ERK/ERK ratios are relative to controls (n=3 retinas for each condition). All values are expressed as the mean ± SEM. Statistics: Mann-Whitney test, *p≤ 0.05, ns: non-significant.

### *Yap^+/-^* mice display retinal dysplasia

We next sought to determine whether *Yap^+/-^* mice exhibit any ocular defect at adult stages when the TAZ-dependent compensatory regulation is no longer effective. Considering the delay in cell cycle exit described above at post-natal stages, and because YAP/TAZ are implicated in the control of organ growth (Yu et al., 2015; Fu et al., 2017), we examined the size of *Yap*^+/-^ eyes. No major difference was found when compared to control eyes (Supplementary Figure S3). Heterozygous *Yap* loss-of-function in humans can result in coloboma (Williamson et al., 2014; Oatts et al., 2017). We, however, did not detect any defects in optic fissure closure in *Yap^+/-^* mouse retina (data not shown). Yet, from P21 onwards, we observed that some mutant retinas displayed one or two dysplastic regions in either the central or dorsal retina (Supplementary Figure S4A, B). The incidence of this non-fully penetrant phenotype was higher in older mice (Supplementary Figure S4C). The severity of the dysplasia is highly variable and each cell layer of the retina is susceptible to be affected (data not shown). Dysplasia were however never detected in the RPE and analysis of the expression of RPE cell markers suggests that the deletion of a *Yap* allele does not disturb RPE integrity (Supplementary Figure S5). Outside the dysplasia, no difference in the thickness of either the outer or the inner nuclear layers was observed between *Yap^+/-^* and control retinas (Supplementary Figure S6). This data indicates that the general structure of the adult retina is not affected in *Yap*^+/-^ mice, apart from the dysplastic region.

### *Yap^+/-^* adult mice display progressive cone photoreceptor degeneration

To assess the impact of one *Yap* allele deletion on adult retinal function, we performed electroretinogram (ERG) recordings in *Yap^+/-^* mice. Scotopic a and b-waves were similar between controls and *Yap^+/-^* mice at all stages examined (Figure 4). This suggests that rod photoreceptors in *Yap^+/-^* mice function normally (reflected by the a-wave), and that visual signal is properly transmitted through the inner retina (reflected by the b-wave). Strikingly however, photopic b-wave amplitude presented a significant depression in 12-month-old *Yap^+/-^* mice compared to controls at high light stimulus intensity (Figure 4). Such specific reduction of the cone-mediated ERG response could signify either defects in the synaptic transmission between cone photoreceptors and bipolar cells and/or the presence of a cone dystrophy. In 12-month-old *Yap^+/-^* retina, IHC analysis showed a significant decrease in the number of Ribeye-positive puncta, which labels the presynaptic ribbons in photoreceptor terminals (Supplementary Figure S7). Ribbons with proper horseshoe shape were present close to dendritic process of the rod-bipolar cell post-synaptic terminals, labelled with anti-Protein Kinase C alpha (PKC-α), suggesting a correct synaptic connection between rod photoreceptor and rod bipolar cells in *Yap^+/-^* retina. In contrast, some ribbons did not exhibit the typical horseshoe shape suggesting compromised synapse integrity (Supplementary Figure S7). We next assessed whether these defects were associated with cone photoreceptor defects in *Yap^+/-^* retina. The number of cones, as inferred by peanut agglutinin (PNA) labelling, was significantly decreased in *Yap^+/-^* ventral retinas compared to controls, affecting both S-opsin and M-opsin labelled cones (Figure 5). S- and M-opsin labelling were indeed both reduced in the ventral retina. S-opsin labelling was also severely decreased in the mid-dorsal (the most dorsal part was not analysed considering the occurrence of dysplasia) and the central retina. In addition to this quantitative phenotype, we also found that the remaining staining was abnormal, being more punctuated compared to the fusiform fluorescence in controls. This phenotype is reminiscent of degenerative cones (Zhang et al., 2017). We confirmed on flat-mounted retinas the punctuated *versus* the fusiform labelling, as well as the reduced number of PNA, S-opsin, and M-opsin labelled cones in the ventral region of 12-month-old *Yap^+/-^* mutant mice (Supplementary Figure S8A). Consistent with a cone degenerative phenotype, we found that Cone Arrestin labelling was also severely decreased in the ventral retina (Supplementary Figure S8B). The cone phenotype was only detected in old mice, as no difference in PNA, S- or M-opsin expression was observed between control and 1- or 4-month-old *Yap*^+/-^ mice (Supplementary Figure S9-S10), consistent with the ERG data. Regarding rods, the distribution of Rhodopsin in 12-month-old mutant and control retinas was similar, showing correct localization within rod outer segments (Supplementary Figure S11). This result also supports the ERG data showing no rod dysfunction. Altogether, these results demonstrate that heterozygous mutation of *Yap* leads to specific cone dystrophy in aged mice.

**Figure 4.**
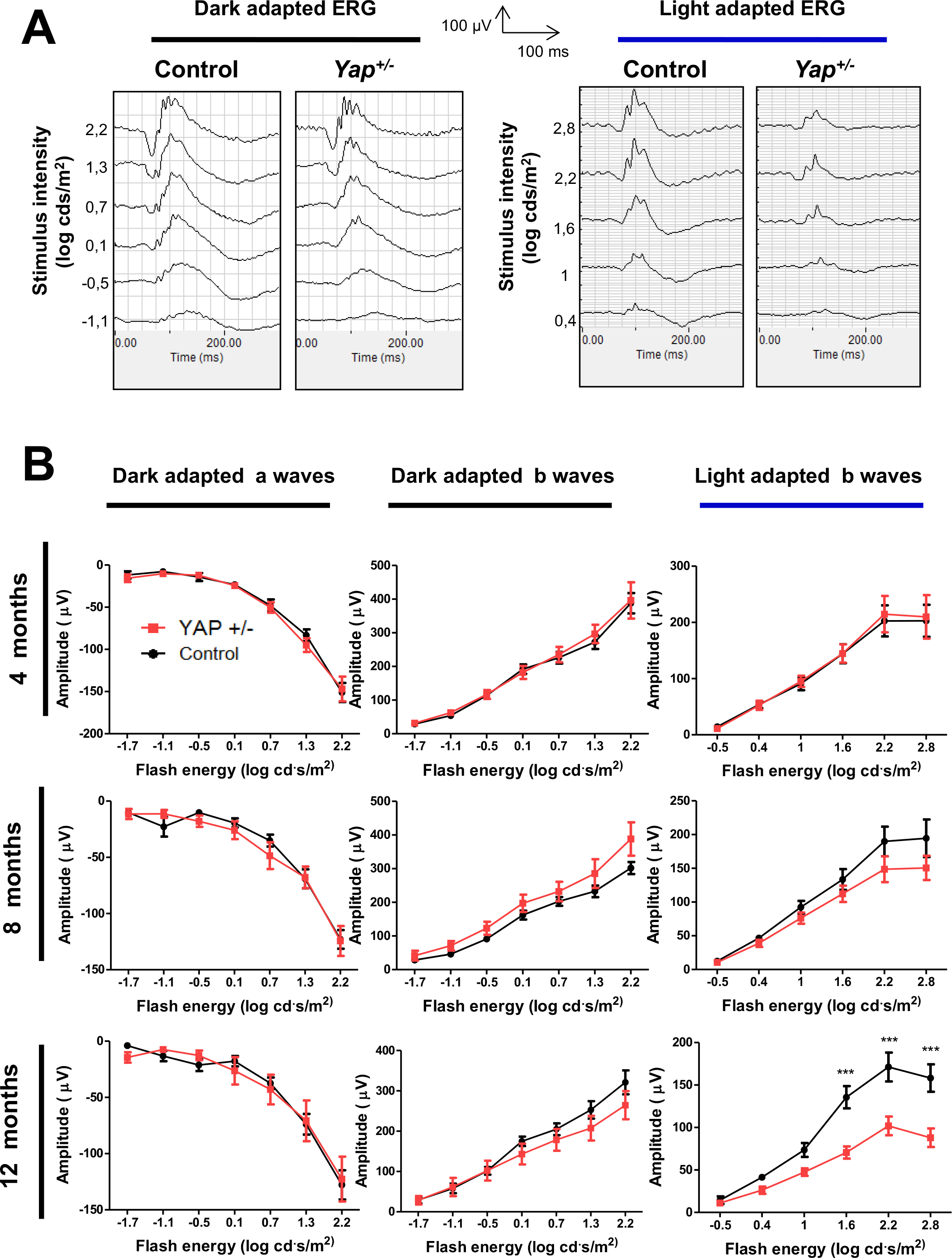
Altered cone-driven vision in *Yap*^+/-^ adult mice. (**A**) Representative ERG intensity series for scotopic (dark-adapted) and photopic (light-adapted) responses in 12-month-old wild-type (Control) and *Yap*^+/-^ mice. (**B**) Quantitative evaluation of the scotopic and photopic a- and b-waves maximum amplitude data from 4-, 8- and 12-month-old wild-type (black) or *Yap*^+/-^ (red) mice. Mean ± SEM intensity response curves are averaged from: 8 controls and 7 *Yap*^+/-^ biological replicates of 4- month-old mice; 9 controls and 8 *Yap*^+/-^ biological replicates of 8-month-old mice; 9 controls and 7 *Yap^+/-^* biological replicates of 12-month-old mice. Statistics: two-way ANOVA test, ***p≤ 0.001.

**Figure 5.**
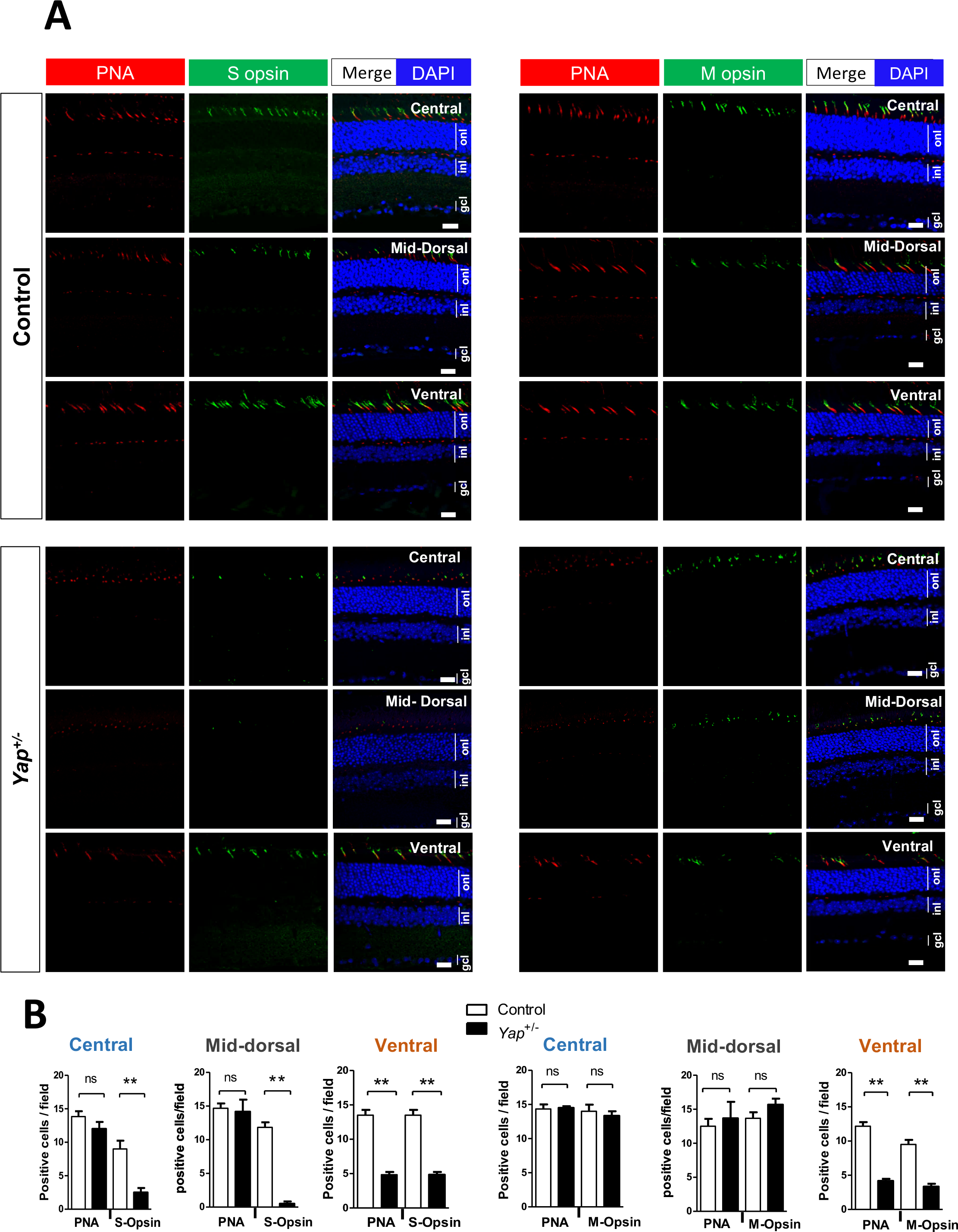
Decreased expression of cone photoreceptor markers in *Yap^+/-^* adult mice. **(A)** Retinal sections from 12-month-old wild type (Control) and *Yap*^+/-^ mice, immunostained for cone makers (PNA, S-Opsin and M-Opsin). Central, mid-dorsal and ventral regions of retinal sections are shown. Nuclei are DAPI counterstained (blue). **(B)** Histograms represent the number of labelled cells per field (150 µm x 150 µm). Values are expressed as the mean ± SEM from at least 3 biological replicates per condition. ONL: outer nuclear layer, INL: inner nuclear layer, GCL: ganglion cell layer. Statistics: Mann-Whitney test, **p≤ 0.01, ns: non-significant. Scale bars: 20 µm.

### Deregulation of genes important for Müller cells homeostasis in *Yap*^+/-^ mice

Since YAP is expressed in adult Müller cells (Hamon et al., 2017) and Müller cells maintain retinal homeostasis (Reichenbach and Bringmann, 2013), we wondered whether cone degeneration in *Yap^+/-^* mice could result from altered Müller cell function. First, we showed that Müller cells were properly located within the inner nuclear layer and that their number was not changed in mutant mice (Supplementary Figure S12). However, RT-qPCR, western blot and IHC analyses revealed that GFAP (glial fibrillary acidic protein) expression, the most sensitive indicator of retinal stress in Müller cells, was dramatically increased in 12-month-old *Yap*^+/-^ retina compared to controls (Figure 6A-C). Therefore, we next explored the impact of *Yap* heterozygous mutation on the expression of Müller cell specific homeostatic regulatory proteins, aquaporin-4 (AQP4) and the potassium channel Kir4.1. Although no significant difference was observed at 8 months, we found that Kir4.1 and AQP4 protein levels were severely reduced in *Yap*^+/-^ mice compared to controls at 12 months of age (Figure 6D, E). Since Müller cells have been shown to actively participate at cone opsins recycling (Mata et al., 2002), we wondered whether the expression of cone specific visual cycle factors could be altered in *Yap^+/-^* Müller cells. We thus analysed the expression of cellular retinaldehyde–binding protein (CRALBP), given its key role in cone visual cycle and its expression in Müller cells (Wang and Kefalov, 2011; Xue et al., 2015). We observed a reduction of CRALPB labelling in both Müller glia and RPE cells of 12-month*-*old *Yap^+/-^* retinas, compared to controls (Figure 7).

**Figure 6.**
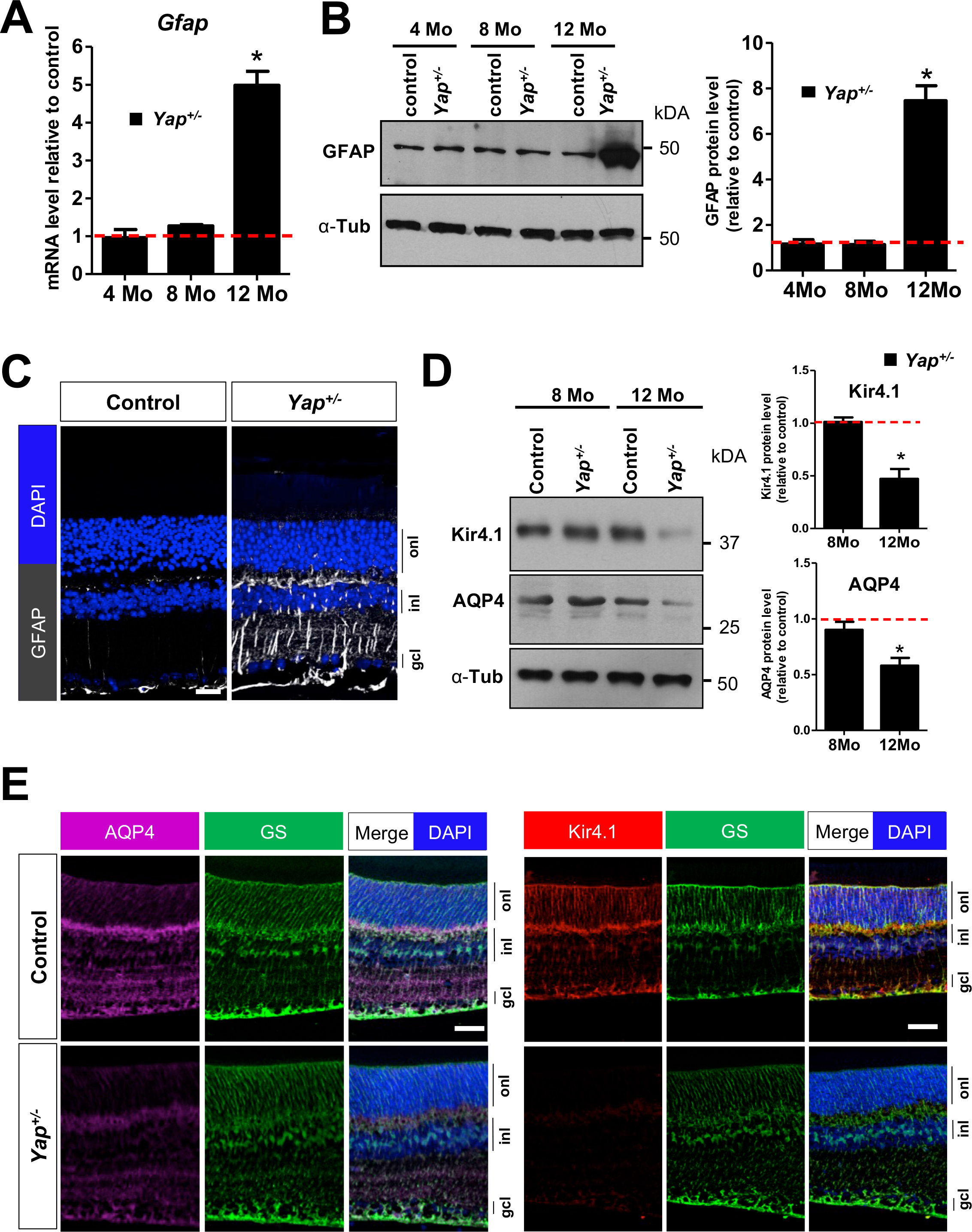
Altered Müller glia homeostasis in *Yap*^+/-^ adult mice. (**A**) RT-qPCR analysis of *Gfap* expression, relative to wild type controls (dashed line). (n= at least 3 biological replicates per condition). (**B**) Analysis of GFAP protein expression level by western blot. Results are normalized to α-tubulin (α-Tub) signal and expressed relative to controls at each stage (dashed line) (n=3 retinas for each condition). (**C**) 12-month-old retinal sections immunostained for GFAP (white). Nuclei are DAPI counterstained. (**D**) Analysis of Kir4.1 and AQP4 protein expression levels by western blot, in 8 or 12-month-old mice. Results are normalized to α-tubulin (α-Tub) signal and expressed relative to controls at each stage (dashed line) (n=3 retinas for each condition). (**E**) 12-month-old retinal sections immunostained for Kir4.1 (red), AQP4 (purple) or the Müller cell marker glutamine synthetase (GS, green). Nuclei are DAPI counterstained (blue). ONL: outer nuclear layer, INL: inner nuclear layer, GCL: ganglion cell layer. All values are expressed as the mean ± SEM. Statistics: Mann-Whitney test, *p≤ 0.05. Scale bar: 50 µm.

**Figure 7.**
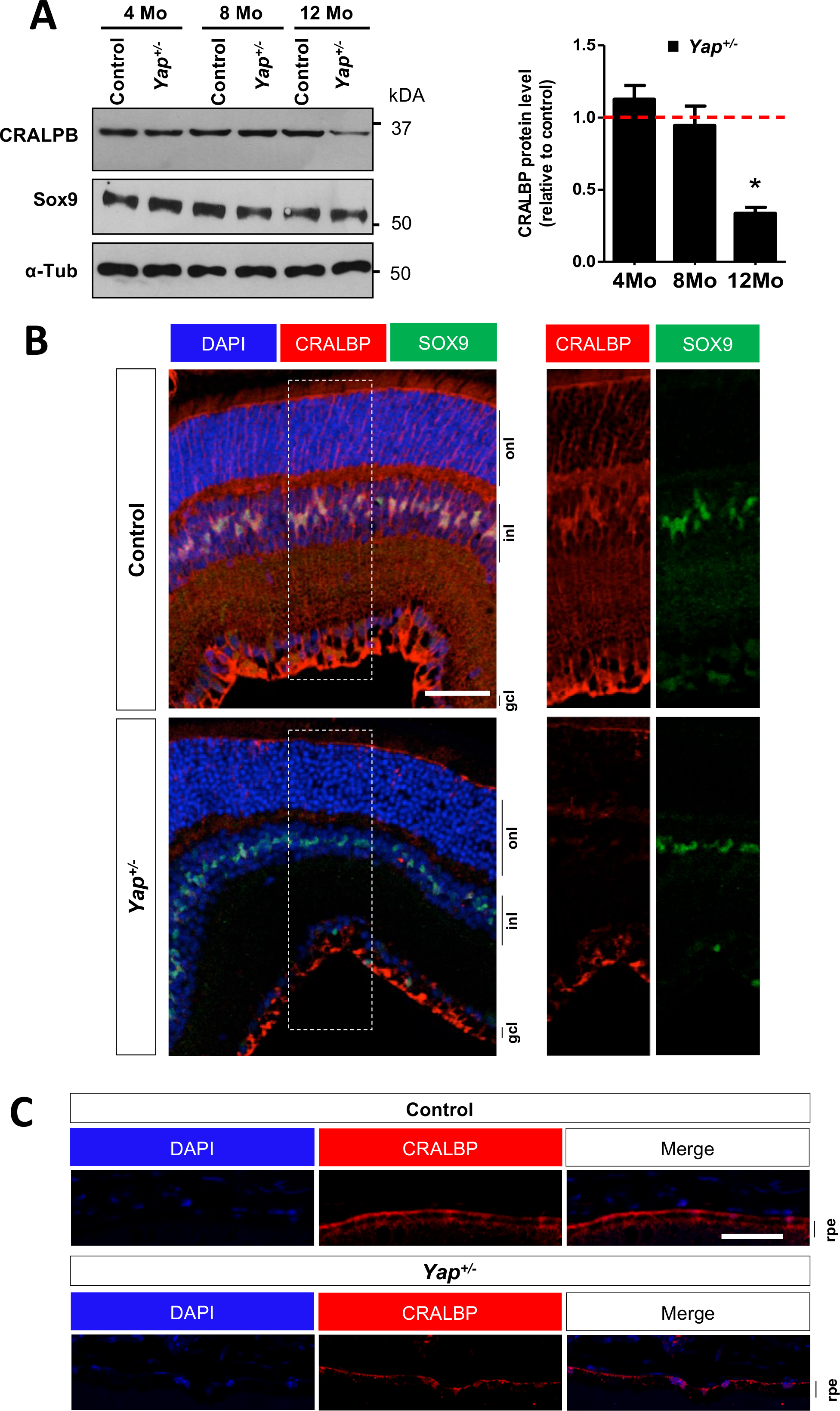
Decreased CRALBP expression in *Yap*^+/-^ adult mice retina. (**A**) Analysis of CRALBP protein expression level by western blot. SOX9 expression serves as a marker of Müller cells. Results are normalized to α-tubulin (α-Tub) signal and expressed relative to controls at each stage (dashed line) (n=3 retinas for each condition). Values are expressed as the mean ± SEM. (**B**) 12-month-old retinal sections immunostained for CRALBP (red) and SOX9 (green). Nuclei are DAPI counterstained (blue). The delineated areas (dashed lines) are enlarged in the right panels. (**C**) 12-month-old RPE sections immunostained for CRALBP (red). Nuclei are DAPI counterstained (blue). INL: inner nuclear layer; ONL: outer nuclear layer; GCL: ganglion cell layer; RPE: retinal pigment epithelium. Statistics: Mann-Whitney test, *p≤ 0.05. Scale bar: 50 µm (B) and 10 µm (C).

Perturbed ionic channels in Müller cells is a leading cause of intraretinal blood vessel defects (Coorey et al., 2012). We thus examined the three retinal vascular plexi in 12-month-old *Yap*^+/-^ mice (Supplementary Figure S13A). We observed a reduction of intermediate vascular plexus without any blood leakage in 12-month-old *Yap*^+/-^ retinal flat-mounts compared to controls (Supplementary Figure S13B and Supplementary Figure S14). Although recent studies have shown that YAP/TAZ are involved in vascular retinal development (Kim et al., 2017), no differences could be observed after labelling the three capillary plexi between 8-month-old *Yap*^+/-^ mice and controls (Supplementary Figure S13B). Such observation ruled out significant developmental defects of vascular networks. Collectively, these data revealed some retinal vasculature defects and altered Müller cell homeostatic function in *Yap^+/-^* aged mice.

### Conditional *Yap* deletion in Müller glia leads to Müller cell homeostasis dysfunction and altered pattern of cone opsin expression

We next sought to investigate whether Müller glia could be the main cell type in which *Yap* haploinsufficiency mediates the observed cone degenerative phenotype. We took advantage of a transgenic line that we previously generated, *Yap^flox/flox^;RaxCreER^T2^*, and that allows Cre-mediated conditional gene ablation specifically in Müller cells (Hamon et al., 2019). It is thereafter named *Yap* CKO while “control” refers to *Yap^flox/flox^* mice. *Yap* deletion was induced in fully differentiated Müller cells, through 4-OHT intraperitoneal injection at P10 (Figure 8A, B). Phenotypic analyses were then conducted on 12-month-old mice. Similar to what we observed in *Yap^+/-^* aged mice, the expression of Müller cell specific homeostatic regulatory proteins AQP4 and Kir4.1 were downregulated and GFAP upregulation indicated reactive gliosis (Figure 8B). Moreover, the expression pattern of S- and M-cone opsins was severely affected compared to control retinas, S-opsin signal being drastically decreased dorsally while M-opsin signal being almost absent ventrally in *Yap* CKO mice (Figure 8C-D). This phenotype is similar to that observed in *Yap^+/-^* retinas. However, the number of cones, as inferred by PNA labelling, was not significantly changed in *Yap* CKO mice compared to controls (Figure 8C,D). Together, although the phenotype thus appears less severe in *Yap* CKO retinas than in *Yap^+/-^* mice, the sole deletion of *Yap* in Müller cell triggers defects in cone opsin expression. These data thus led us to propose a model in which YAP function in Müller cells would be necessary for ensuring the maintenance of Müller cell homeostasis for preserving cone integrity (Figure 9).

**Figure 8.**
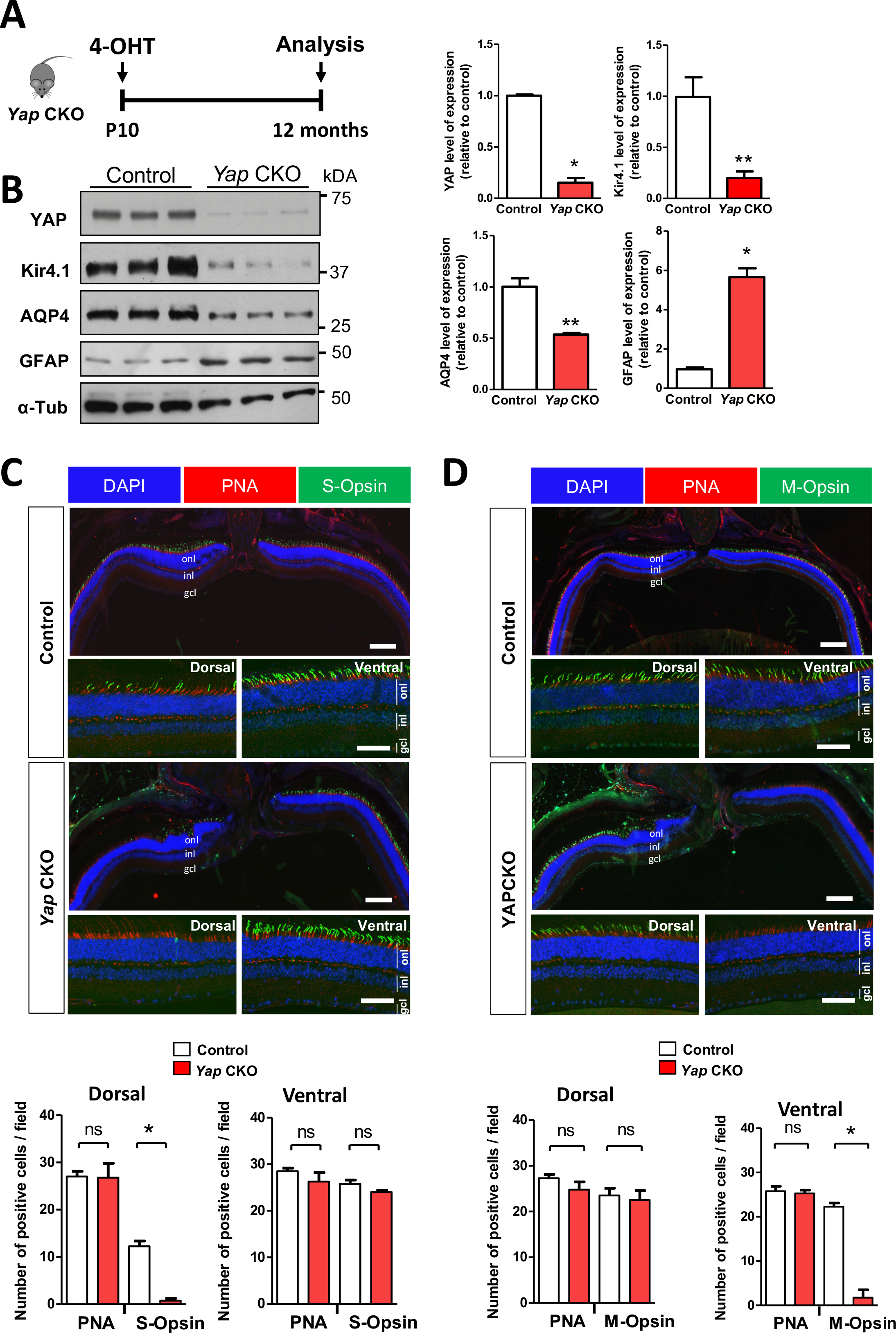
Müller cell dysfunction and decreased expression of S/M-opsin in *Yap* CKO. (**A**) Timeline diagram of the experimental procedure used in B. *Yap^flox/flox^* mice (control) or *Yap^flox/flox^;Rax-CreER^T2^* mice (*Yap CKO*) received a single dose of 4-OHT at P10 and their retinas were analyzed at 12 months. (**B**) Analysis of the protein expression level of AQP4, Kir4.1, and GFAP. Results are normalized to α-tubulin (α-Tub) signal and expressed relative to control (n=3 retinas for each condition). (**C-D**) Retinal sections from 12-month-old control and *Yap CKO* mice immunostained for S-opsin (green) and PNA (red) (C) or M-opsin (green) and PNA (red) (D). Dorsal, central and ventral sections of retinal sections are shown. Nuclei are DAPI counterstained (blue). Histograms represent the number of labelled cells per field (250 µm x 250 µm). Values are expressed as the mean ± SEM from at least 3 biological replicates per condition. ONL: outer nuclear layer, INL: inner nuclear layer, GCL: ganglion cell layer. Statistics: Mann-Whitney test, *p≤ 0.05, **p≤ 0.01, ns: non-significant. Scale bars: 100 µm (central part) (enlarged panels) and 50 µm (dorsal and ventral part).

## DISCUSSION

While YAP function during eye development has been well documented, our data demonstrate the usefulness of *Yap* heterozygous mice to better understand YAP function in the adult retina. Interestingly, our study revealed that compensatory upregulation of YAP partner, TAZ, could not be sustained throughout life and declined during aging. This dynamic regulatory mechanism results in *Yap* haploinsufficiency only in the aged retina. We propose a model where the maintenance of proliferative progenitor cells in *Yap^+/-^* post-natal retina would result from the compensatory TAZ activity, leading to a gain of function-like phenotype, mediated by EGFR pathway potentiation (Figure 9). During aging, when this compensatory mechanism is no longer operational in *Yap*^+/-^ mice, we discovered the occurrence of an age-related cone dystrophy, associated with reduced cone-driven vision. Our data suggest that this phenotype may be the consequence of impaired Müller cell homeostatic function. This study therefore reveals that deregulation of *Yap* gene dosage could cause retinal degenerative diseases.

**Figure 9.**
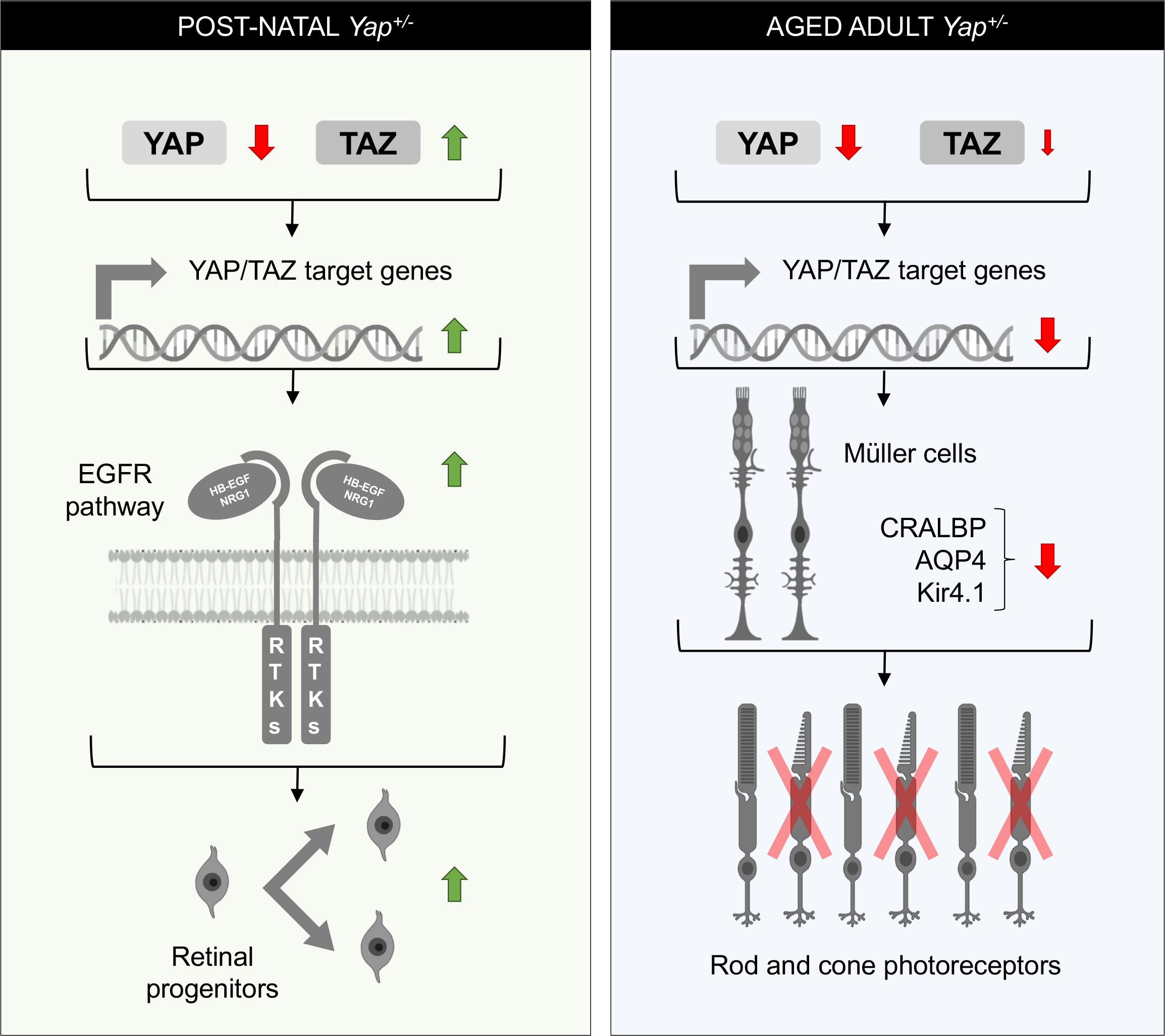
Proposed model explaining *Yap*^+/-^ phenotypes at post-natal and adult stages. Our results revealed the occurrence of a TAZ compensatory mechanism in post-natal *Yap*^+/-^ retina, *i.e.* the decreased of *Yap* expression (red arrow) is accompanied by an increase of *Taz* expression (green arrow), that results in the upregulation of YAP/TAZ target gene expression. Our data suggest that this leads to prolonged proliferation of post-natal retinal progenitor cells, and that this phenotype may likely result from EGFR pathway potentiation. In the adult *Yap*^+/-^ retina, the compensatory mechanism is no longer effective, and YAP/TAZ target genes expression is diminished. Under such conditions, one *Yap* allele deletion progressively leads to impaired Müller cell homoeostasis. We propose that this may, at least in part, contribute to impaired cone-specific visual cycle and therefore to the cone dystrophy observed in aged *Yap*^+/-^ mice. RTKs: Receptor tyrosine kinases. This figure was created with some schemas from ©BioRender - biorender.com.

It has previously been reported that changes in YAP abundance results in compensatory regulation of TAZ to maintain Hippo signaling homeostasis (Finch-Edmondson et al., 2015). We interestingly found that one *Yap* allele deletion was indeed accompanied by an increase in *Taz* expression in the retina, as well as an increase in *Tead* gene expression. However, such rises appeared to go beyond *Yap* compensation as we observed an increase in YAP/TAZ target genes. Interestingly, this dosage compensation system was not maintained in the retina of adult mice after 4 months of age. All these quantitative data on the abundance of YAP/TAZ and their target genes highlight *Yap^+/-^* mice as an excellent model to study the role of YAP in the adult retina, when TAZ no longer compensates for *Yap* deletion. Unlike in humans, where heterozygous loss-of-function mutations in *YAP* causes coloboma (Williamson et al., 2014; Oatts et al., 2017), *Yap*^+/-^ mice do not exhibit such congenital malformation of the eye. A plausible hypothesis is that the compensatory mechanism during eye development is not regulated in humans as it is in mice. Interestingly, the phenotype in humans is not fully penetrant, which could indicate potential YAP/TAZ dosage variations between individuals.

YAP overexpression in neonatal mouse retinas was shown to promote cell proliferation and inhibit cell cycle exit of late retinal progenitors (Zhang et al., 2012). We also found a delay in cell cycle exit in *Yap^+/-^* post-natal retina. We propose that this is the consequence of the compensatory regulation that leads to TAZ level increase. Indeed, despite the genetic *Yap^+/-^* context, we observed an upregulation of YAP/TAZ target genes at post-natal stages. Furthermore, our data suggest that this delayed cell cycle exit of retinal progenitor may be mediated by an increase in EGFR activity. Previous studies in other cell types have demonstrated that YAP and TAZ promote proliferation by potentiating the EGFR pathway (He et al., 2015; Chen and Harris, 2016; Yang et al., 2016). This is also consistent with our recent finding suggesting that YAP is a key regulator of the EGFR pathway in adult reactive Müller cells in a degenerative context (Hamon et al., 2019). It remains to be investigated whether EGFR pathway components are directly regulated by YAP/TAZ in retinal cells. We raised the hypothesis that the prolonged maintenance of cell proliferation at post-natal stages could be linked to the appearance of dysplastic regions in *Yap^+/-^* retina. Indeed, an excessive number of retinal cells may subsequently lead to abnormal folding of retinal layers and trigger such dysplasia at later stages. Alternatively, considering that YAP regulates key effectors of matrix stiffening (Dupont Exp cell research 2016), we can also hypothesize that the stiffness of the retinal tissue is affected, leading to retinal folding. This could underlie the increased incidence of dysplasia observed during aging.

We showed previously that YAP is expressed in Müller cells in the adult retina (Hamon et al., 2017). We thus raised the hypothesis that cone degeneration in *Yap^+/-^* mice could result from altered Müller cell function. Indeed, Müller cells play many essential roles in retinal maintenance, regulation of synaptic activity, ion and water homeostasis, and importantly in mediating cone visual cycle (Reichenbach and Bringmann, 2013). Moreover, conditional Müller cell ablation was shown to cause photoreceptor degeneration (Shen et al., 2012; Byrne et al., 2013). Our hypothesis is supported by the finding that deleting *Yap* specifically in Müller cells in *Yap* CKO mice also leads to abnormal expression of cone opsins. Both mutants exhibit reactive gliosis and impaired expression of genes involved in Müller cell homeostatic function (AQP4 and Kir4.1) consistent with a model where impaired cone integrity would be caused by Müller cell dysfunction.

We also discovered in *Yap^+/-^* Müller glia an altered expression of CRALBP, which is involved in cone-specific visual cycle. Knock-down or mutations in CRALBP gene leads to decreased cone-driven ERG responses in zebrafish, M-cone loss in mice or can lead to severe cone photoreceptor-mediated retinal disease in patients (Maw et al., 1997; Fleisch et al., 2008; Xue et al., 2015). Importantly, restoration of CRALBP expression specifically in Müller cells, but not RPE cells, can rescue the sensitivity of CRALBP-deficient cones in the mouse (Xue et al., 2015). We can thus raise the hypothesis that the severe decrease of CRALBP levels in Müller cells of aged *Yap^+/-^* mice may contribute to the observed cone dysfunction and degeneration. We did not observe any significant decrease in CRALBP levels in *Yap* CKO retinas (data not shown), which may explain, at least in part, the less severe cone phenotype (PNA staining unaffected) compared to the one observed in *Yap^+/-^* mice. Besides, the decreased abundance of YAP in *Yap^+/-^* mice in other cell types than Müller cells may also explain the different severity of the phenotype. Although we did not detect any major defects in RPE cells in *Yap^+/-^* mice, we observed a decreased expression of CRALBP in RPE cells. We therefore cannot exclude the possibility that the RPE is not functioning properly, contributing to cone degeneration. Similarly, the defect in the intermediate vascular plexus observed in *Yap^+/-^* mice could also participate to the degenerative phenotype.

We here did not focus our analysis on extraocular tissues but we did not detect any major cornea or lens defects (data not shown). On the other hand, ocular phenotypes in a *Yap^+/-^* mouse line were recently examined with an emphasis on the cornea and severe corneal pathology was reported (Kim et al., 2019). In this mutant mouse, similar to our findings, retinal defects were detected only in aged mice. However, in contrast to our study, the phenotype was very severe, including retinal detachment, defect in the RPE and important loss of photoreceptors. It remains to be addressed whether genetic background differences could explain these phenotypic variabilities between the two *Yap^+/-^* mouse lines.

To conclude, we highlighted in this study a novel role for YAP in the maintenance of cone photoreceptors in adult mice, associated with a regulation of Müller cell homeostasis. Overall, our data showing cone degeneration in *Yap^+/-^* mice (*i*) warrant further investigation in patients with *YAP* heterozygous loss-of-function mutations, regarding their cone-driven vision during aging, and (*ii*) identify *Yap* as a novel candidate gene that could account for cone dystrophy in patients with unidentified mutations.

## ACKNOWLEDGEMENTS

We are thankful to S. Meilhac, Jeff Wrana, and Seth Blackshaw for mouse lines. We are grateful to S. Lourdel for her help with the maintenance of mouse colonies. This research was supported by grants to M.P. from the FRM (Fondation pour la Recherche Médicale), Association Retina France, Fondation Valentin Haüy, DIM Région Île de France and UNADEV. DGG is a FRM fellow.

---

**Supplementary Figure S1.**
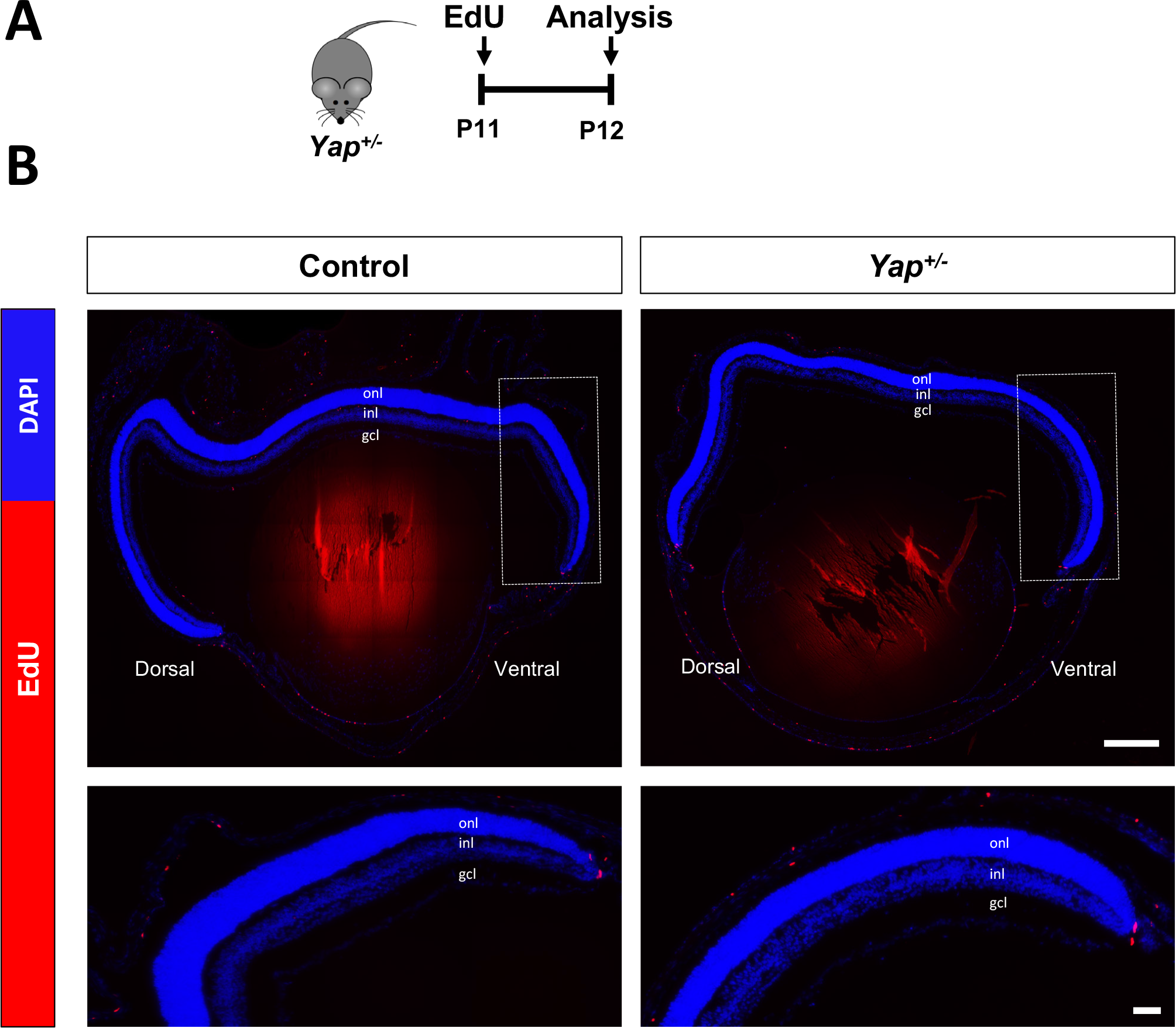
Absence of retinal progenitor proliferation in P12 *Yap^+/-^* mice. (**A**) Timeline diagram of the experimental procedure used in B. Wild-type (Control) or *Yap^+/-^* mice were injected with EdU at P11 and the presence of EdU-positive cells was assessed 24 hours later (P12). (**B**) Retinal sections labelled for EdU (red) and DAPI counterstained (blue). Ventral areas are enlarged in the bottom panels. INL: inner nuclear layer; ONL: outer nuclear layer; GCL: ganglion cell layer. Scale bar: 200 µm and 50 µm (enlarged panels).

**Supplementary Figure S2.**
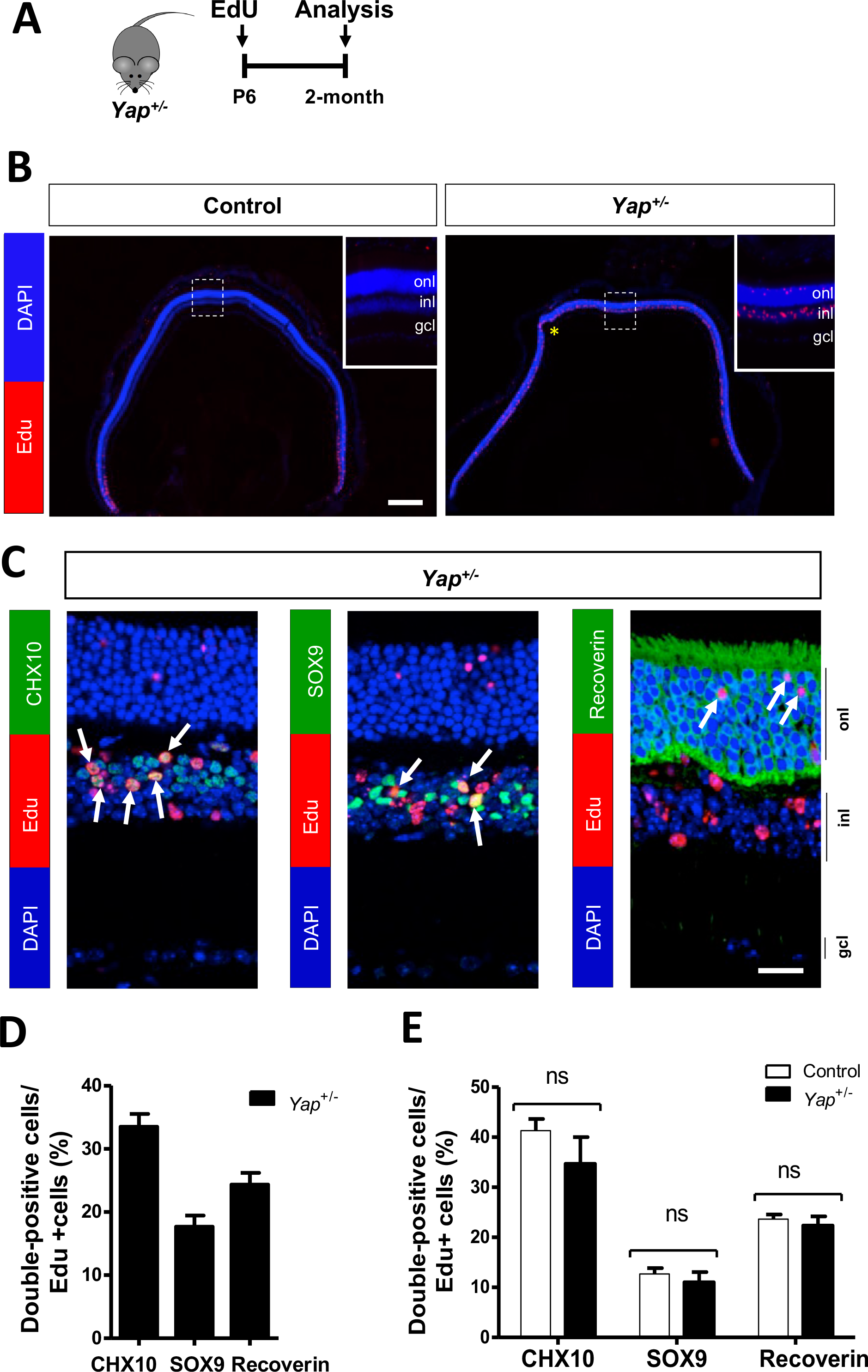
Correct cell fate of retinal progenitors with delayed cell cycle exit in *Yap^+/-^* mice. (**A**) Timeline diagram of the birthdating experimental procedure used in B. Wild-type (Control) or *Yap^+/-^* mice were injected with EdU at P6 and the fate of EdU-positive cells was analysed in 2-month-old mice. (**B**) Retinal sections labelled for EdU (red) and DAPI counterstained (blue). The delineated areas are enlarged in the insets. The asterisk marks the location of a dysplastic region. (**C**) Retinal sections labelled for EdU (red) and stained for the indicated markers (SOX9, CHX10, Recoverin, green) and DAPI counterstained (blue). Arrows point to double labelled cells. (**D, E**) Quantification of double labelled cells among EdU-positive cells per field (400 µm x 400 µm) in the central (D) or peripheral (E) retinal region. Mean values ± SEM from 3 retinas per condition are shown. INL: inner nuclear layer; ONL: outer nuclear layer; GCL: ganglion cell layer. Statistics: Mann-Whitney test, ns: non-significant. Scale bar: 200 µm (B), 50 µm (C).

**Supplementary Figure S3.**
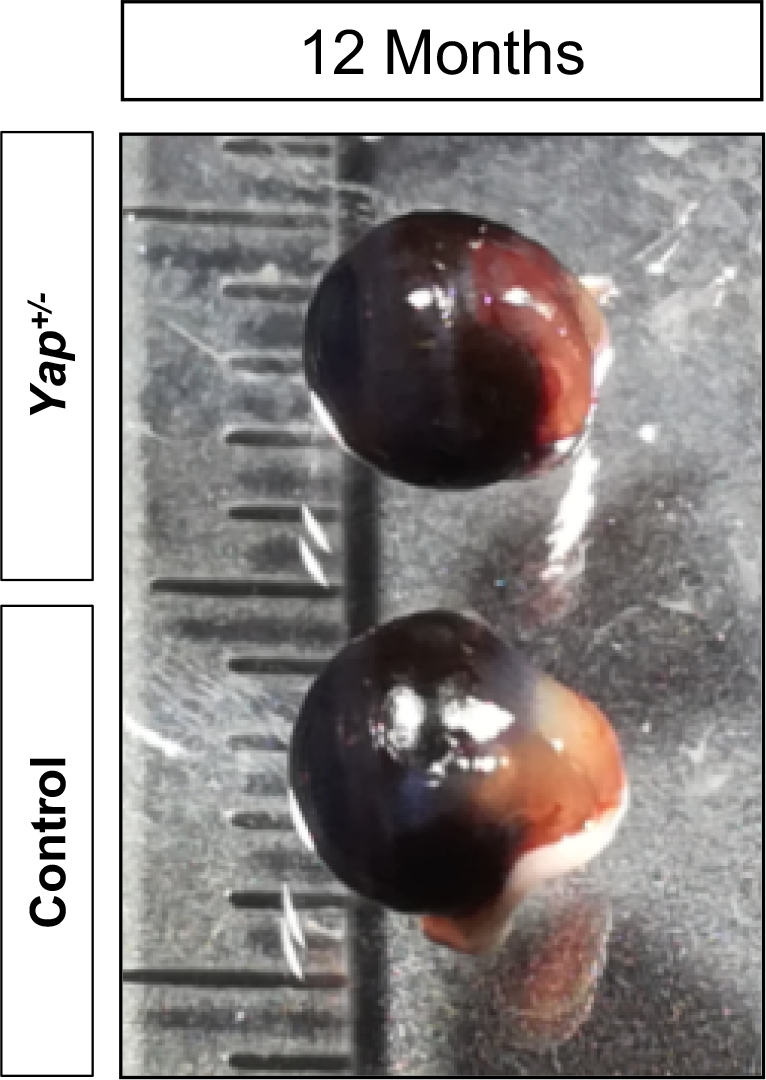
One allele deletion of *Yap* does not alter eye size. Enucleated eyes from WT and *Yap*^+/-^ 12-month-old mice.

**Supplementary Figure S4.**
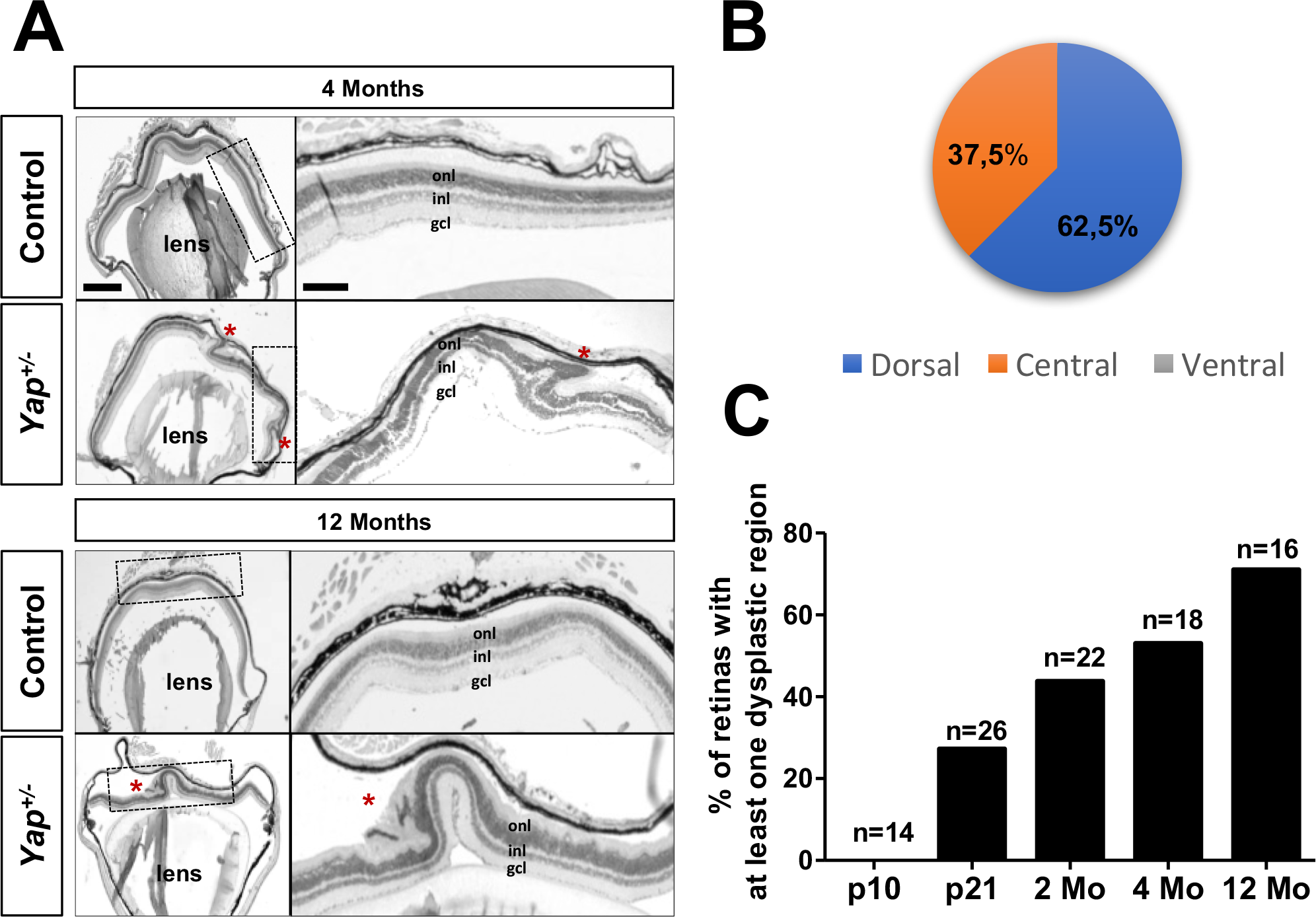
Retinal dysplasia in *Yap^+/-^* mice. (**A**) H&E staining of retinal sections in 4 and 12-month-old wild type (Control) and *Yap^+/-^* mice. Red asterisks indicate the position of dysplastic regions. Area delineated with dashed black lines are enlarged in panels on the right showing finger-like protrusions of *Yap^+/-^* retinal layers toward the RPE. (**B**) Number of dorsal, central or ventral retinal dysplasia in *Yap*^+/-^ retinas expressed in percentage of the total number of dysplasia (n=96 retinas). (**C**) Incidence of retinal dysplasia in control and *Yap*^+/-^ mice at different stages. INL: inner nuclear layer; ONL: outer nuclear layer; GCL: ganglion cell layer. Scale bar: 200 µm and 50 µm (enlarged panels).

**Supplementary Figure S5.**
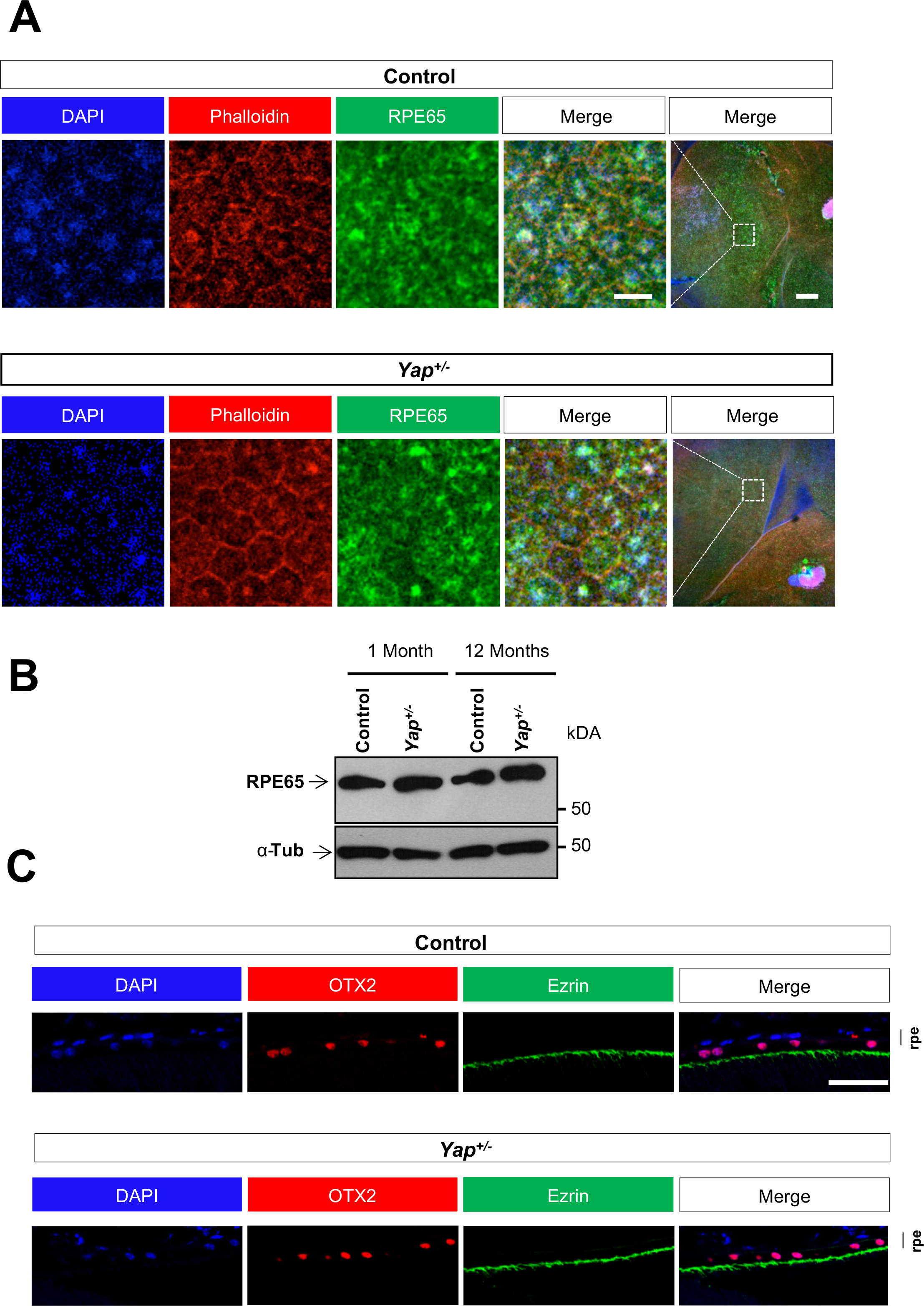
Normal appearance of the RPE in *Yap^+/-^* mice. (**A**) RPE flat mounts from 12-month-old wild type (control) and *Yap^+/-^* mice immunostained for Phalloidin (red) and RPE65 (green). Nuclei are DAPI counterstained (blue). (**B**) Analysis of RPE65 and α-tubulin (α-Tub) level of expression in 1 or 12-month-old mice. (**C**) RPE sections from 12-month-old mice immunostained for OTX2 (red) and Ezrin (green). Nuclei are DAPI counterstained (blue). RPE: retinal pigment epithelium. Scale bar: 200 µm and 20 µm (enlarged panels) (A), 10 µm (C).

**Supplementary Figure S6.**
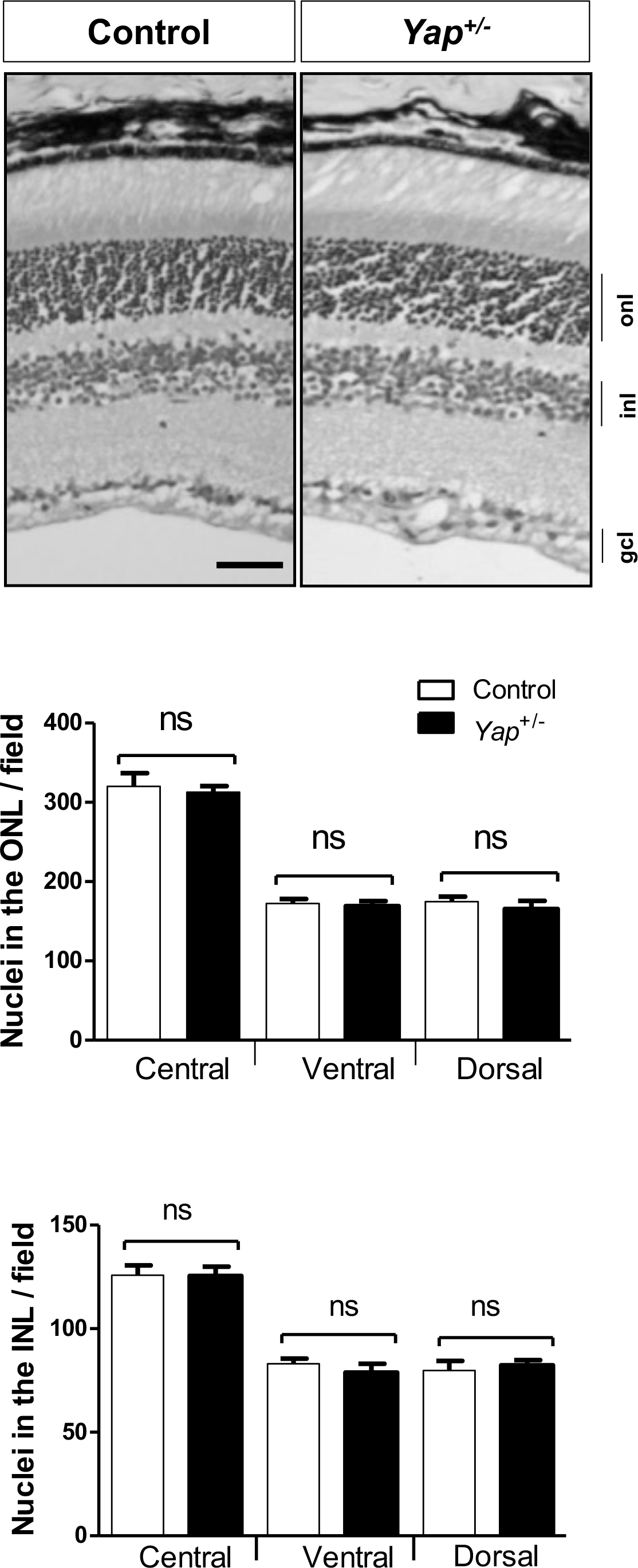
Normal thickness of *Yap^+/-^* adult retina. H&E staining of retinal sections from 12-month-old wild type (Control) and *Yap^+/-^* mice. A region away from a dysplastic region is shown. Histogram represents the measurement of the outer and inner nuclear layer thickness. Mean values ± SEM from three independent retinas are shown. INL: inner nuclear layer; ONL: outer nuclear layer; GCL: ganglion cell layer. Statistics: Mann-Whitney test, ns: non-significant. Scale bar: 50 µm.

**Supplementary Figure S7.**
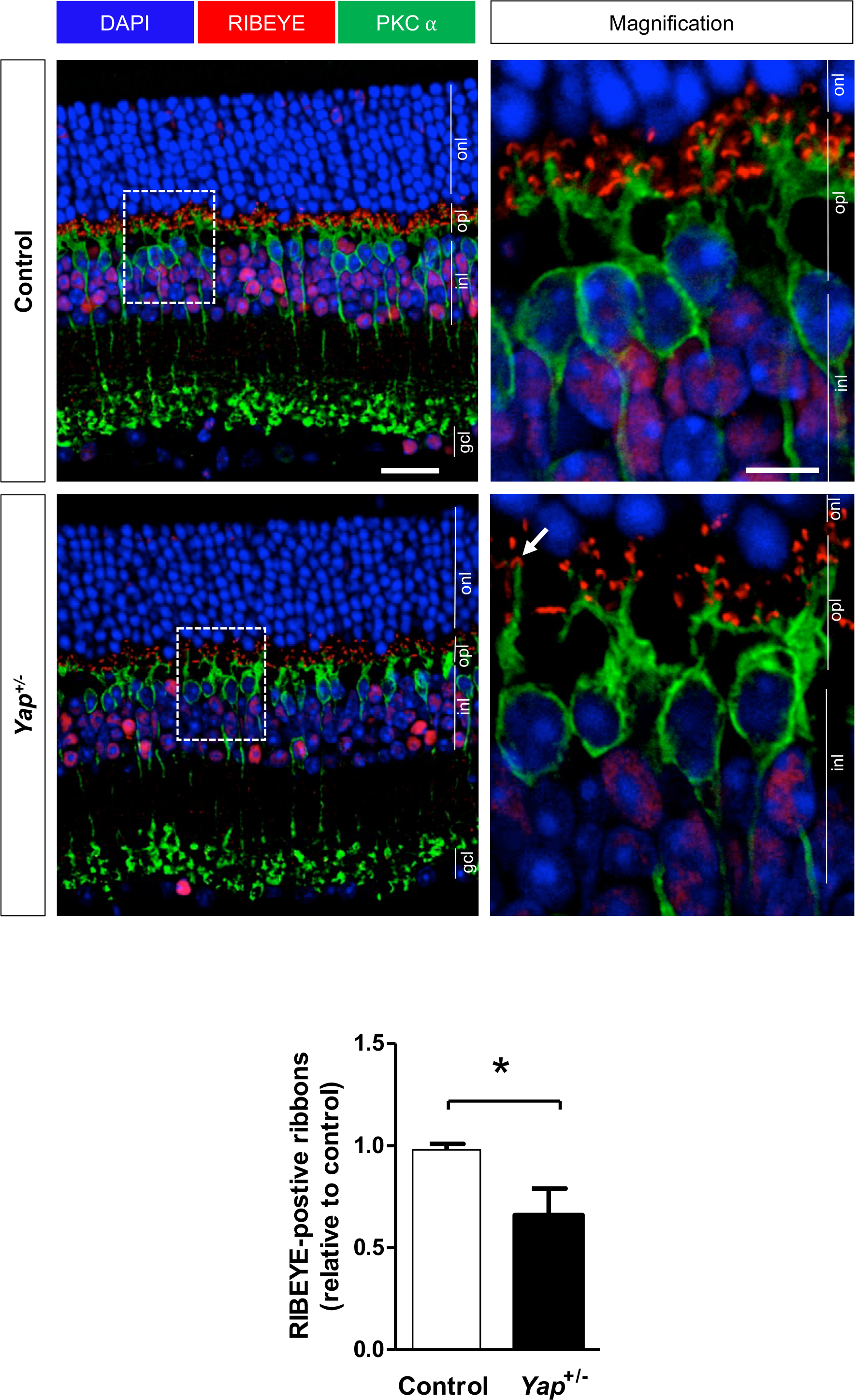
Altered architecture of the photoreceptor ribbon synapse in *Yap*^+/-^ adult mice. 12-month-old retinal sections immunostained for PKC-*α* (green) and RIBEYE (red). Delineated areas (dashed lines) are enlarged in the right panels. Nuclei are DAPI counterstained (blue). An arrow indicates a ribbon with a normal horseshoe shape, facing PKC-α labelled rod-bipolar cell post-synaptic terminals in *Yap^+/-^* retinas. The histogram indicates the number of labelled RIBEYE positive ribbons per field (150 µm x 150 µm). ONL: outer nuclear layer, INL: inner nuclear layer, GCL: ganglion cell layer. Scale bar: 20 µm and 50 µm (enlarged panels).

**Supplementary Figure S8.**
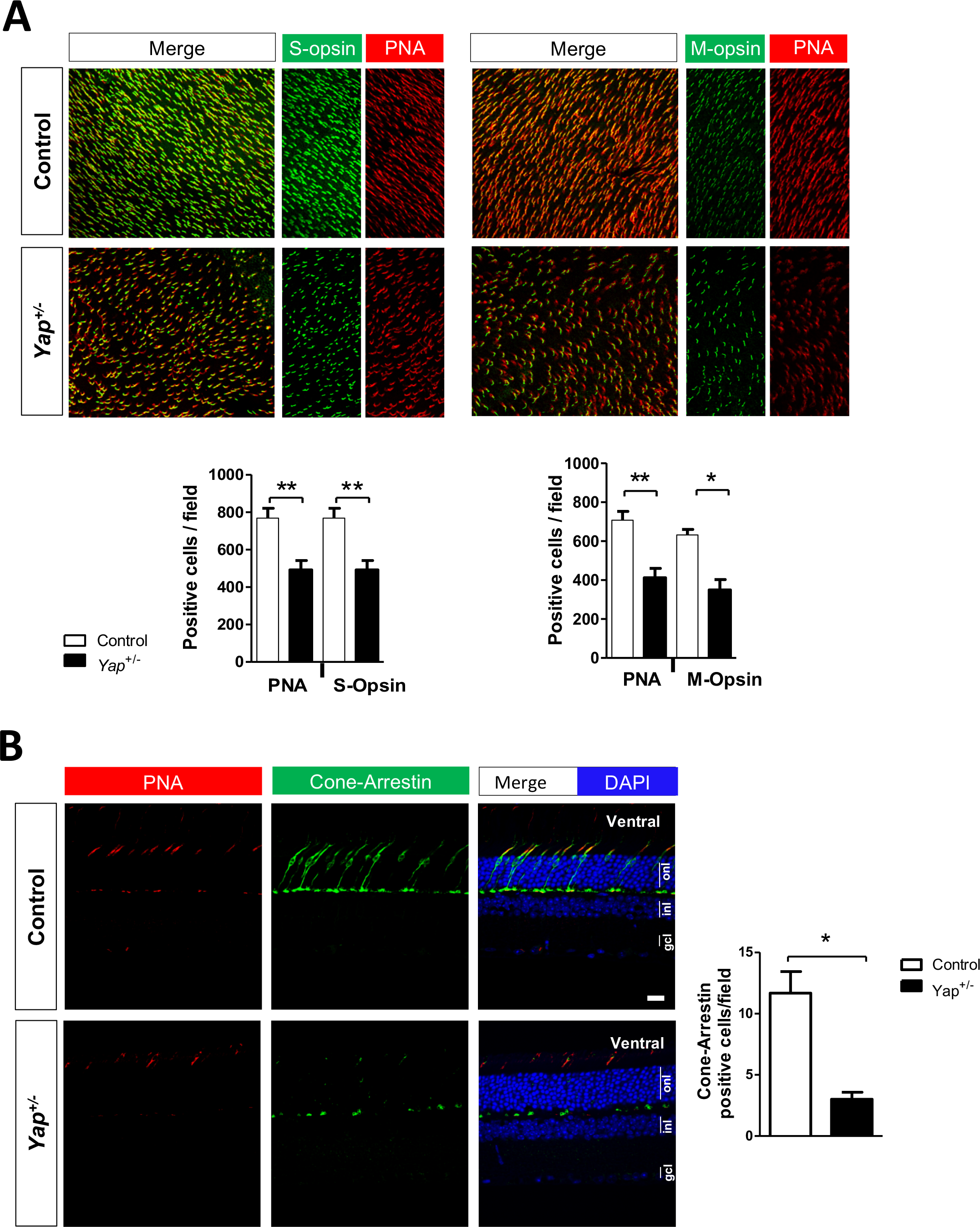
Decreased expression of cone markers in *Yap*^+/-^ mice retinas. **(A)** Retinal flat-mounts from 12-month-old wild type (control) and *Yap*^+/-^ mice, immunostained for cone makers (PNA, S-Opsin and M-Opsin) in the ventral part of the retina. Histograms indicate the number of labelled cells per field (400 µm x 400 µm). **(B)** 12-month-old mouse retinal sections immunostained for PNA (red) and Cone Arrestin (green). Nuclei are DAPI counterstained (blue). Histograms represent the number of Cone Arrestin labelled cells per field (150 µm x 150 µm). Values are expressed as the mean ± SEM from at least 3 biological replicates per condition. ONL: outer nuclear layer, INL: inner nuclear layer, GCL: ganglion cell layer. Statistics: Mann-Whitney test, *p≤ 0.05, **p≤ 0.01.

**Supplementary Figure S9.**
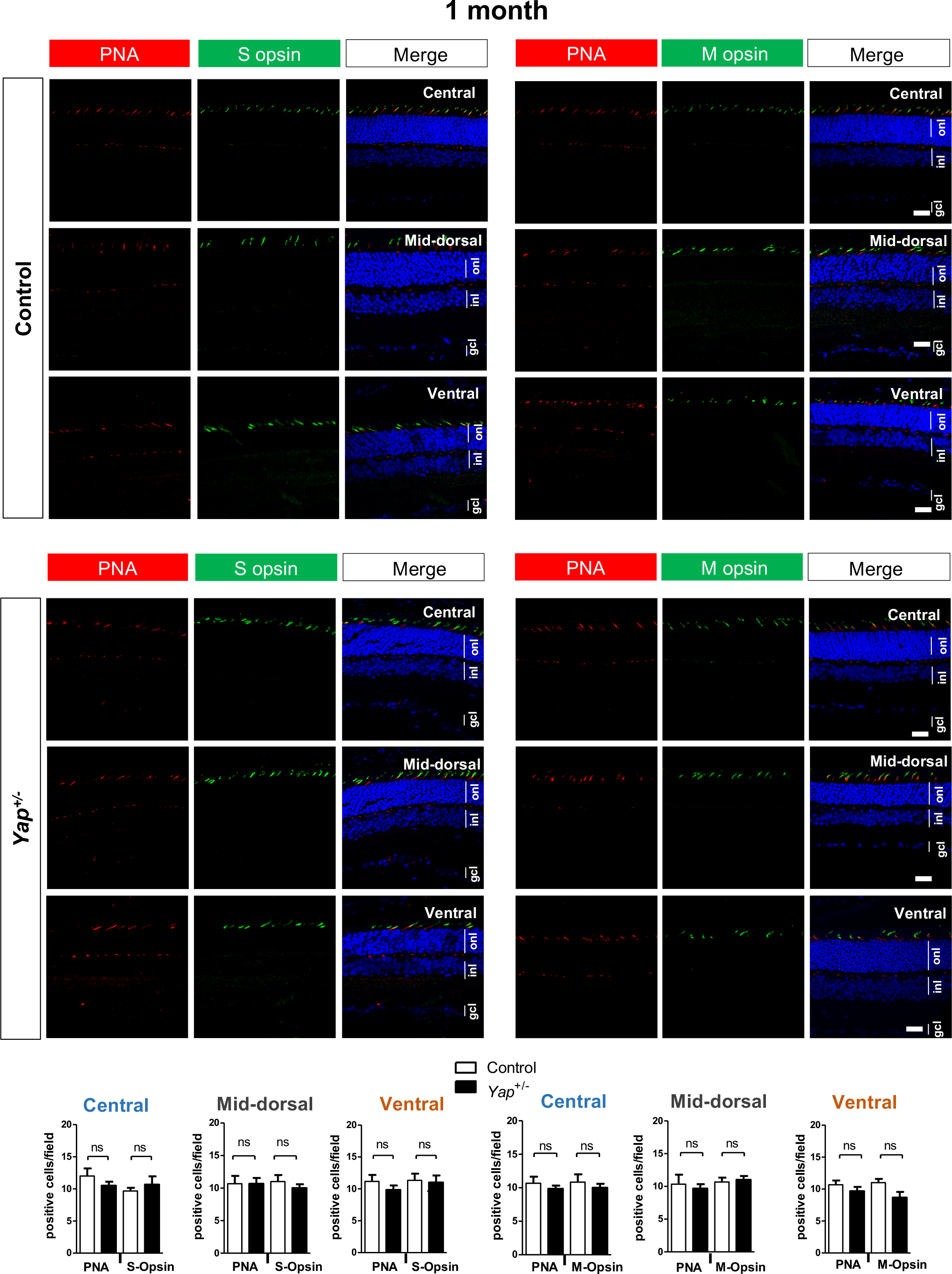
Normal expression of cone markers in 1-month-old *Yap*^+/-^ mice. 1-month-old retinal sections immunostained for cone markers (PNA, S-opsin, M-opsin). Histograms represent the quantification of labelled cells per field (400 µm x 400 µm). Central, mid-dorsal and ventral regions of retinal sections are shown. Mean values ± SEM from 3 retinas per condition are shown. Statistics: Mann-Whitney test, ns: non-significant. INL: inner nuclear layer; ONL: outer nuclear layer; GCL: ganglion cell layer. Scale bar: 20 µm.

**Supplementary Figure S10.**
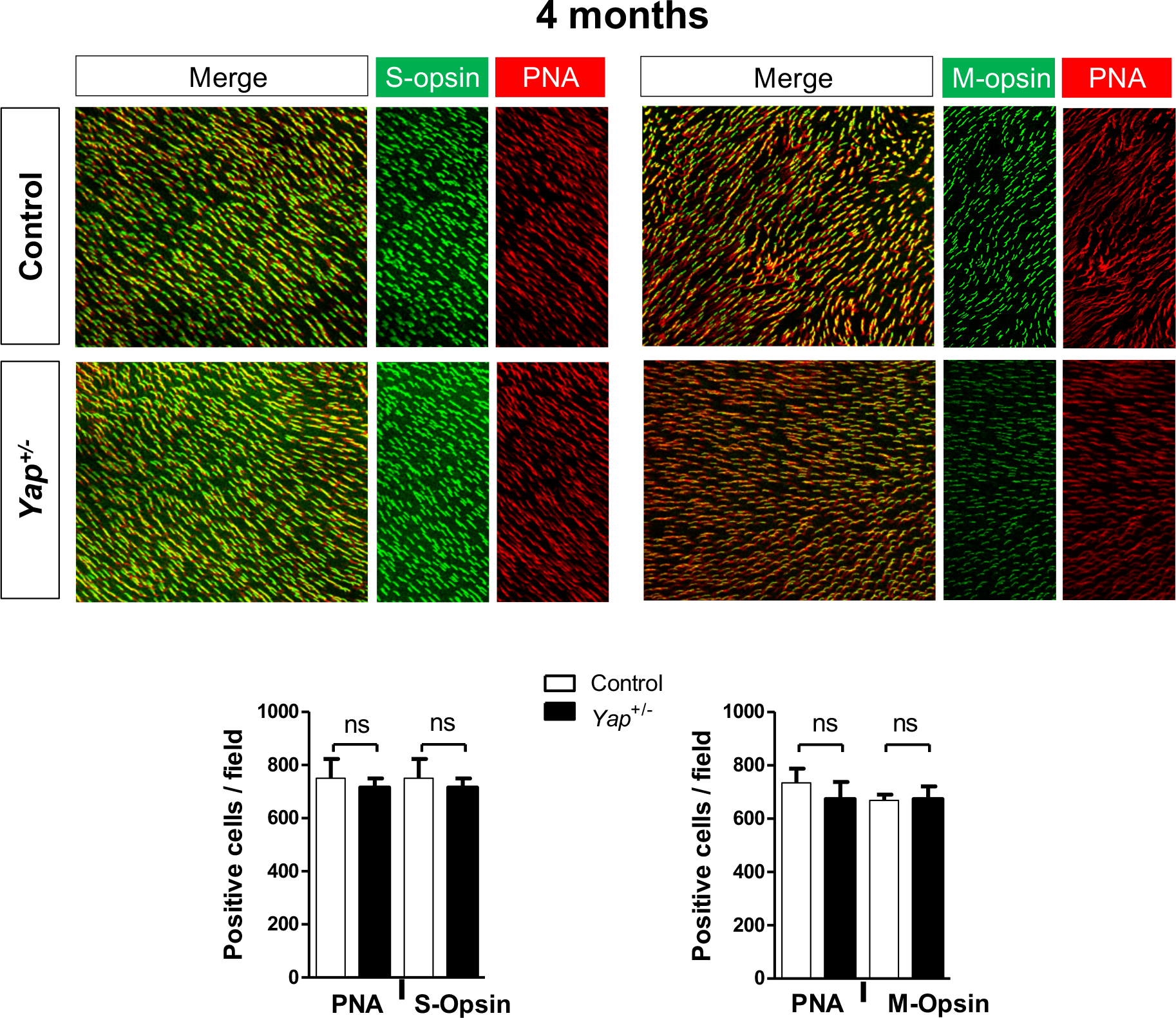
Normal expression of cone markers in 4-month-old *Yap*^+/-^ mice. 4-month-old retinal flat-mounts immunostained for cone markers (PNA, S-opsin, M-opsin). Histograms represent the quantification of labelled cells per field (150 µm x 150 µm). Mean values ± SEM from 3 retinas per condition are shown. Statistics: Mann-Whitney test, ns: non-significant.

**Supplementary Figure S11.**
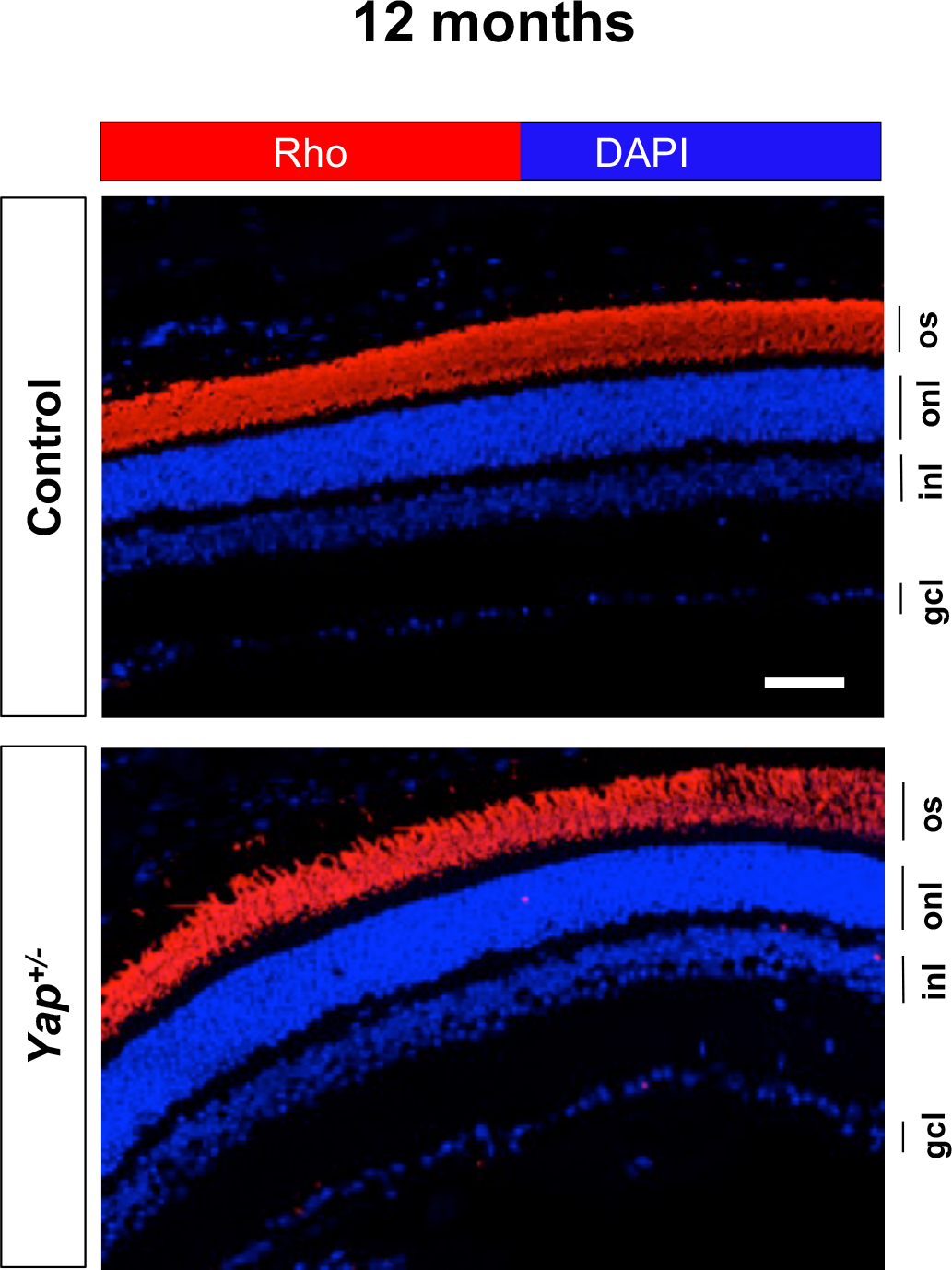
Normal expression of Rhodopsin in *Yap*^+/-^ adult mice. 12-month-old retinal sections immunostaining for Rhodopsin (Rho, red). Nuclei are DAPI counterstained (blue). OS: outer segment, INL: inner nuclear layer; ONL: outer nuclear layer; GCL: ganglion cell layer. Scale bar: 50 µm.

**Supplementary Figure S12.**
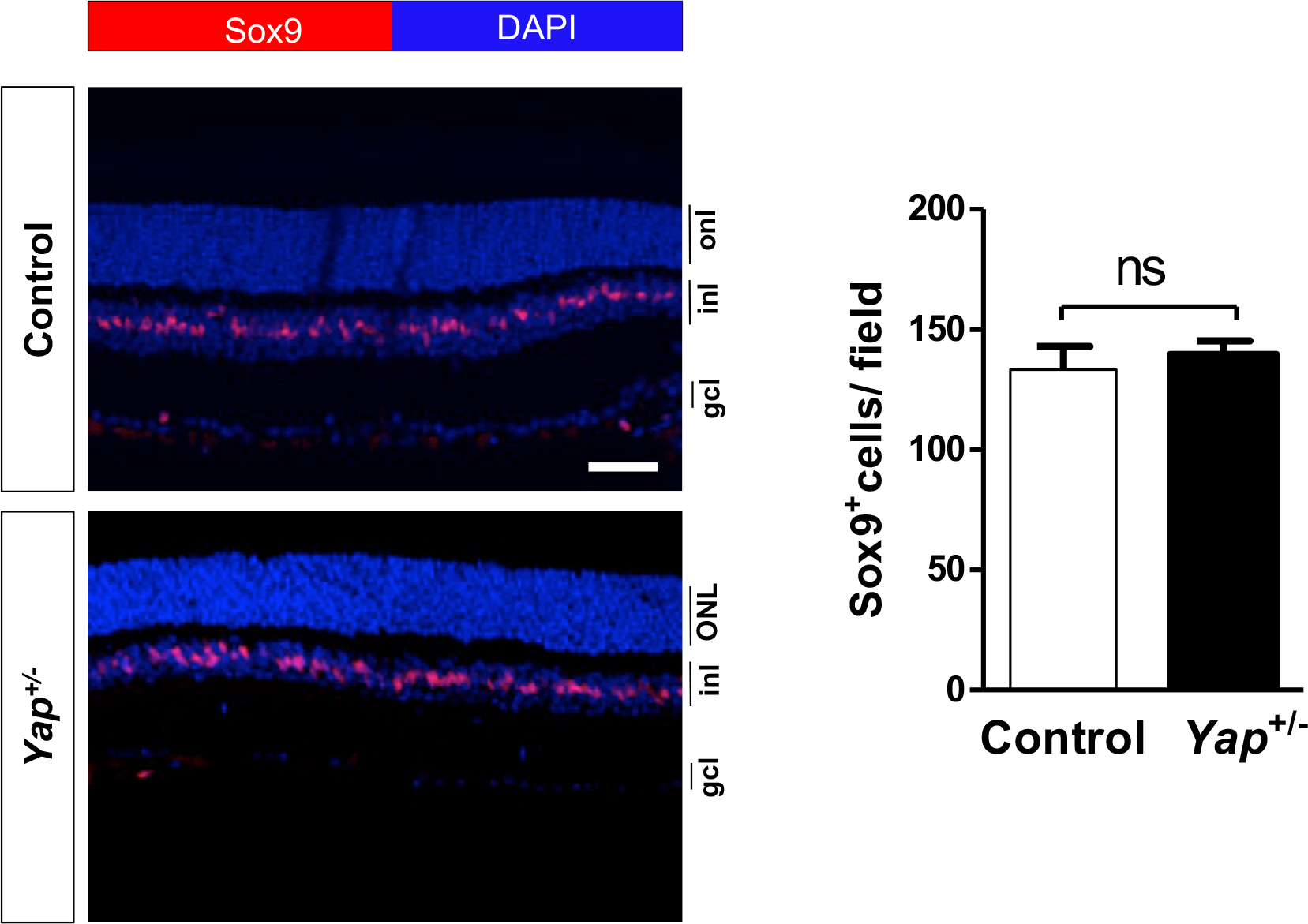
Normal number of Müller cells in *Yap*^+/-^ adult mice. 2-month-old retinal sections immunostained for SOX9 (green) and counterstained with DAPI to identify the nuclei (blue). Histogram represents the quantification of labelled cells per field (1200 µm x 1200 µm). Mean values ± SEM from 3 retinas per condition are shown. INL: inner nuclear layer; ONL: outer nuclear layer; GCL: ganglion cell layer. Statistics: Mann-Whitney test, ns: non-significant. Scale bar: 50 µm.

**Supplementary Figure S13.**
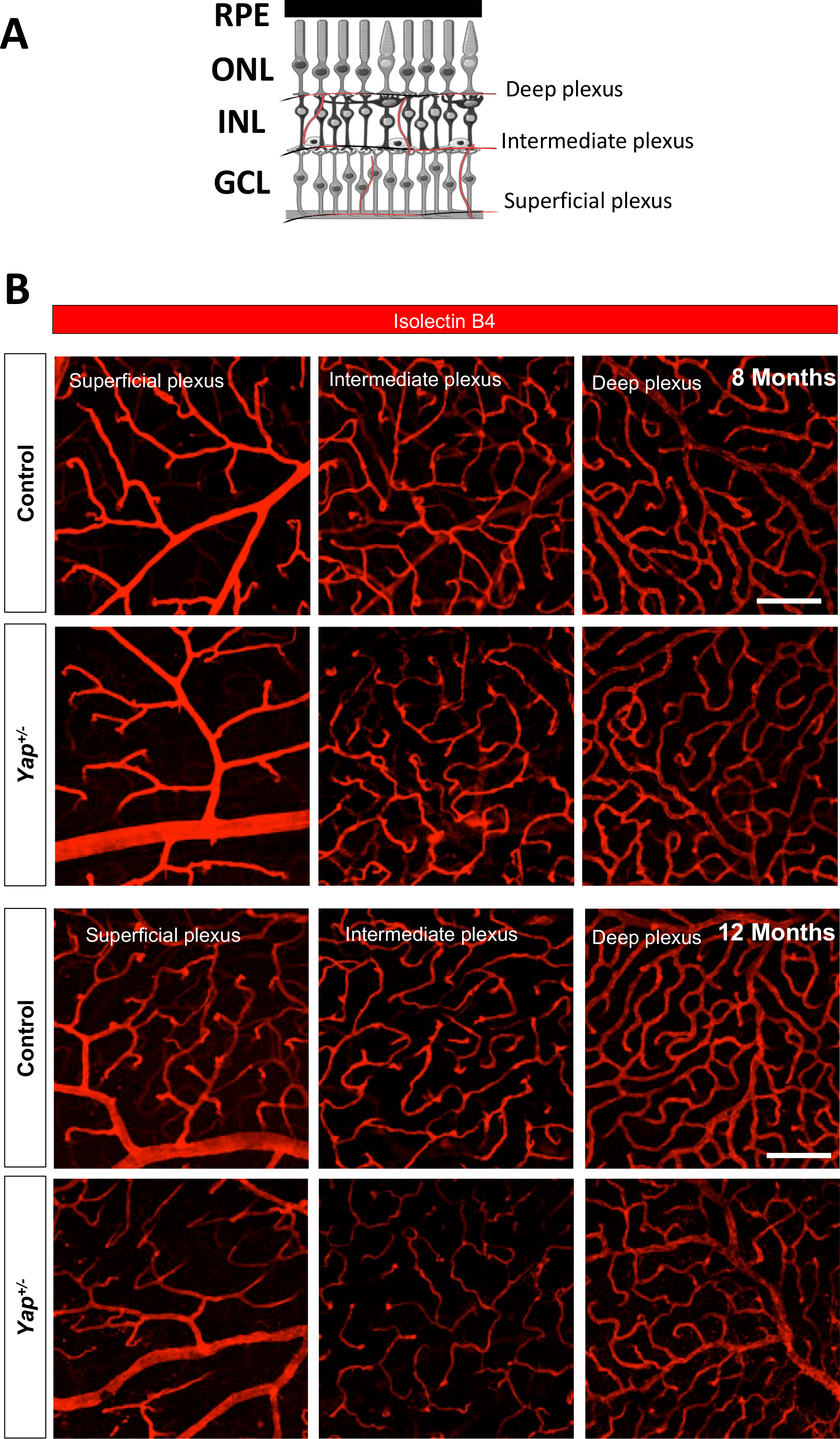
Alteration of intermediate retinal vascular plexus in *Yap*^+/-^ adult mice. (**A**) Schematic representation of the three-retinal plexi: deep plexus, intermediate plexus and superficial plexus (red) in the mouse retina. This figure was created with some schemas adapted from Servier Medical Art by Servier (licensed under a Creative Commons Attribution 3.0 Unported License). (**B**) 8 and 12-old-month retinal flat-mounts stained for endothelial cells with isolectin B_4_ (red) in the superficial, intermediate, and deep plexi. INL: inner nuclear layer; ONL: outer nuclear layer; GCL: ganglion cell layer; RPE: retinal pigment epithelium. Scale bar: 100 µm.

**Supplementary Figure S14.**
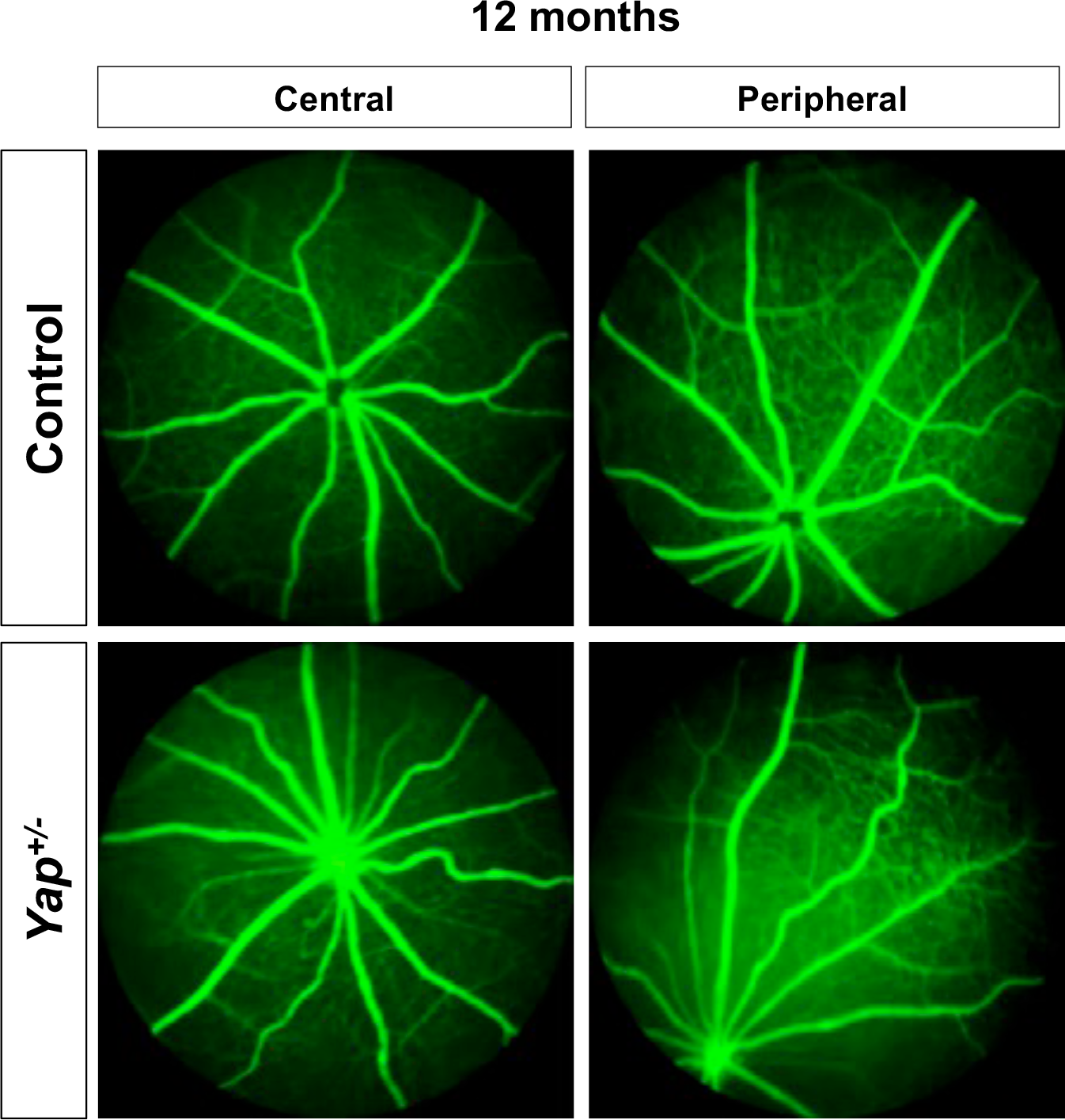
No blood leakage in *Yap*^+/-^ adult mice. Fluorescein angiography in 12-old-month mouse retina showing no signs of vascular leakage approximately 2 min after intra-peritoneal injection of sodium fluorescein. Scale bar: 200 µm.

**Supplementary Table S1:**
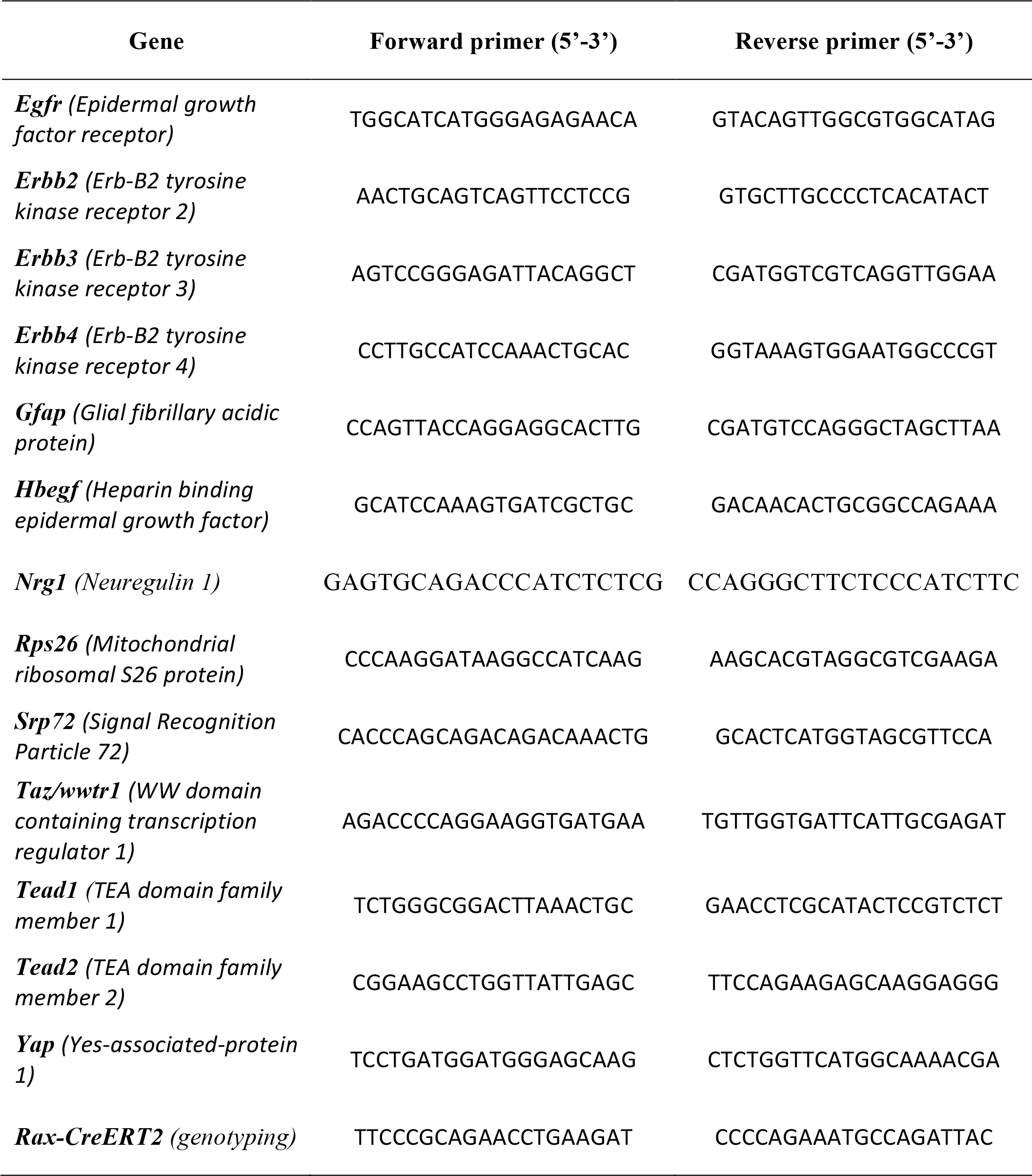
List of primers.

**Supplementary Table S2:**
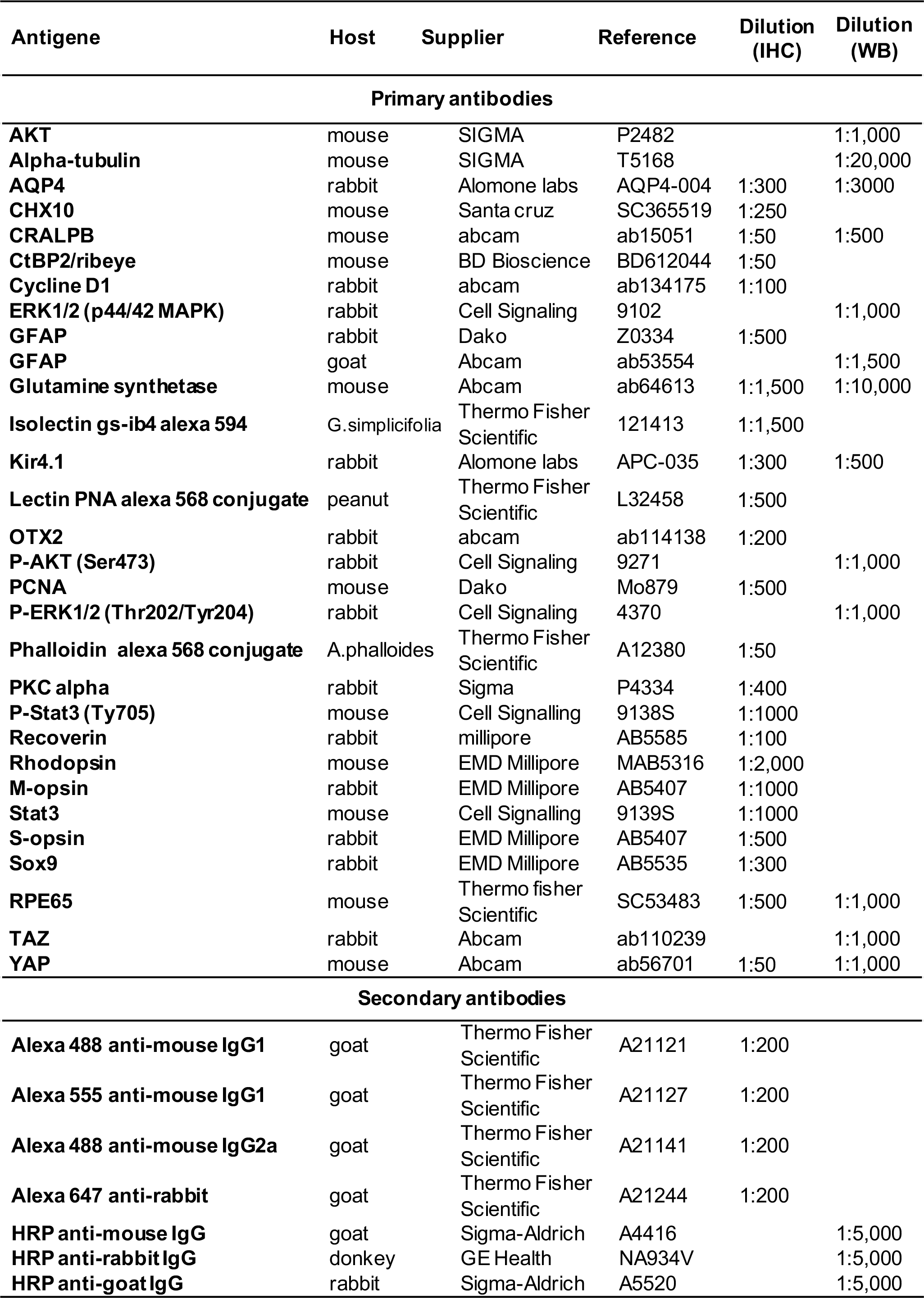
List of antibodies. IHC: immunohistochemistry, WB: western blot.

| Antigene | Host | Supplier | Reference | Dilution (IHC) | Dilution (WB) |
| --- | --- | --- | --- | --- | --- |
| **Primary antibodies** |  |  |  |  |  |
| **AKT** | mouse | SIGMA | P2482 |  | 1:1,000 |
| **Alpha-tubulin** | mouse | SIGMA | T5168 |  | 1:20,000 |
| **AQP4** | rabbit | Alomone labs | AQP4-004 | 1:300 | 1:3000 |
| **CHX10** | mouse | Santa cruz | SC365519 | 1:250 |  |
| **CRALPB** | mouse | abcam | ab15051 | 1:50 | 1:500 |
| **CtBP2/ribeye** | mouse | BD Bioscience | BD612044 | 1:50 |  |
| **Cycline D1** | rabbit | abcam | ab134175 | 1:100 |  |
| **ERK1/2 (p44/42 MAPK)** | rabbit | Cell Signaling | 9102 |  | 1:1,000 |
| **GFAP** | rabbit | Dako | Z0334 | 1:500 |  |
| **GFAP** | goat | Abcam | ab53554 |  | 1:1,500 |
| **Glutamine synthetase** | mouse | Abcam | ab64613 | 1:1,500 | 1:10,000 |
| **Isolectin gs-ib4 alexa 594** | G.simplicifolia | Thermo Fisher Scientific | 121413 | 1:1,500 |  |
| **Kir4.1** | rabbit | Alomone labs | APC-035 | 1:300 | 1:500 |
| **Lectin PNA alexa 568 conjugate** | peanut | Thermo Fisher Scientific | L32458 | 1:500 |  |
| **OTX2** | rabbit | abcam | ab114138 | 1:200 |  |
| **P-AKT (Ser473)** | rabbit | Cell Signaling | 9271 |  | 1:1,000 |
| **PCNA** | mouse | Dako | Mo879 | 1:500 |  |
| **P-ERK1/2 (Thr202/Tyr204)** | rabbit | Cell Signaling | 4370 |  | 1:1,000 |
| **Phalloidin alexa 568 conjugate** | A.phalloides | Thermo Fisher Scientific | A12380 | 1:50 |  |
| **PKC alpha** | rabbit | Sigma | P4334 | 1:400 |  |
| **P-Stat3 (Ty705)** | mouse | Cell Signalling | 9138S | 1:1000 |  |
| **Recoverin** | rabbit | millipore | AB5585 | 1:100 |  |
| **Rhodopsin** | mouse | EMD Millipore | MAB5316 | 1:2,000 |  |
| **M-opsin** | rabbit | EMD Millipore | AB5407 | 1:1000 |  |
| **Stat3** | mouse | Cell Signalling | 9139S | 1:1000 |  |
| **S-opsin** | rabbit | EMD Millipore | AB5407 | 1:500 |  |
| **Sox9** | rabbit | EMD Millipore | AB5535 | 1:300 |  |
| **RPE65** | mouse | Thermo fisher Scientific | SC53483 | 1:500 | 1:1,000 |
| **TAZ** | rabbit | Abcam | ab110239 |  | 1:1,000 |
| **YAP** | mouse | Abcam | ab56701 | 1:50 | 1:1,000 |
| **Secondary antibodies** |  |  |  |  |  |
| **Alexa 488 anti-mouse IgG1** | goat | Thermo Fisher Scientific | A21121 | 1:200 |  |
| **Alexa 555 anti-mouse IgG1** | goat | Thermo Fisher Scientific | A21127 | 1:200 |  |
| **Alexa 488 anti-mouse IgG2a** | goat | Thermo Fisher Scientific | A21141 | 1:200 |  |
| **Alexa 647 anti-rabbit** | goat | Thermo Fisher Scientific | A21244 | 1:200 |  |
| **HRP anti-mouse IgG** | goat | Sigma-Aldrich | A4416 |  | 1:5,000 |
| **HRP anti-rabbit IgG** | donkey | GE Health | NA934V |  | 1:5,000 |
| **HRP anti-goat IgG** | rabbit | Sigma-Aldrich | A5520 |  | 1:5,000 |

